# Clamp loader processing is important during DNA replication stress

**DOI:** 10.1101/2022.11.10.516041

**Authors:** Tommy F. Tashjian, Peter Chien

**Author notes:** Peter Chien, Dept. of Biochemistry and Molecular Biology, University of Massachusetts Amherst, Life Science Laboratories Room N325, 240 Thatcher Rd, Amherst MA 01003-9364, (413)545-2310, @chienlab.

## Abstract

The DNA clamp loader is critical to the processivity of the DNA polymerase and coordinating synthesis on the leading and lagging strands. In bacteria the major subunit of the clamp loader, DnaX, has two forms: the essential full-length τ and shorter γ. These are conserved across bacterial species and three distinct mechanisms have been found to create them: ribosomal frameshift, transcriptional slippage, and, in *Caulobacter crescentus*, proteolysis. This conservation suggests that DnaX processing is evolutionarily important, but its role remains unknown.

Here we find a bias against switching from expression of a wild type *dnaX* to a nonprocessable *τ-only* allele in *Caulobacter*. Despite this bias, cells are able to adapt to the *τ-only* allele with little effect on growth or morphology and only minor defects during DNA damage. Motivated by transposon sequencing, we find that loss of the gene *sidA* in the *τ-only* strain slows growth and increases filamentation. Even in the absence of exogenous DNA damage treatment, the *ΔsidA τ-only* double mutant shows induction of and dependance on *recA*, likely due to a defect in resolution of DNA replication fork stalling. We find that some of the phenotypes of the *ΔsidA τ-only* can be complemented by expression of γ but that an overabundance of τ-only *dnaX* is also detrimental. The data presented here suggest that DnaX processing is important during resolution of replication fork stalling events during DNA replication stress.

**IMPORTANCE:** Though the presence of DnaX τ and γ forms is conserved across bacteria, different species have developed different mechanisms to make these forms. This conservation and independent evolution of mechanisms suggest that having two forms of DnaX is important. Despite having been discovered more than 30 years ago, the purpose of expressing both τ and γ is still unclear. Here, we present evidence that expressing two forms of DnaX and controlling the abundance and/or ratio of the forms is important during the resolution of replication fork stalling.

## INTRODUCTION

The clamp loader is an essential component of the DNA replication machinery. It performs two critical functions: coordinating DNA synthesis on the leading and lagging strands (1) and loading the DNA clamp (2, 3) that increases the processivity of the DNA polymerase. There is also evidence that the clamp loader modulates the access to the replication fork of factors important for DNA damage bypass (4), DNA repair (5–7), DNA replication termination (8, 9), and replication restart (10) (Figure 1).

**Figure 1.**
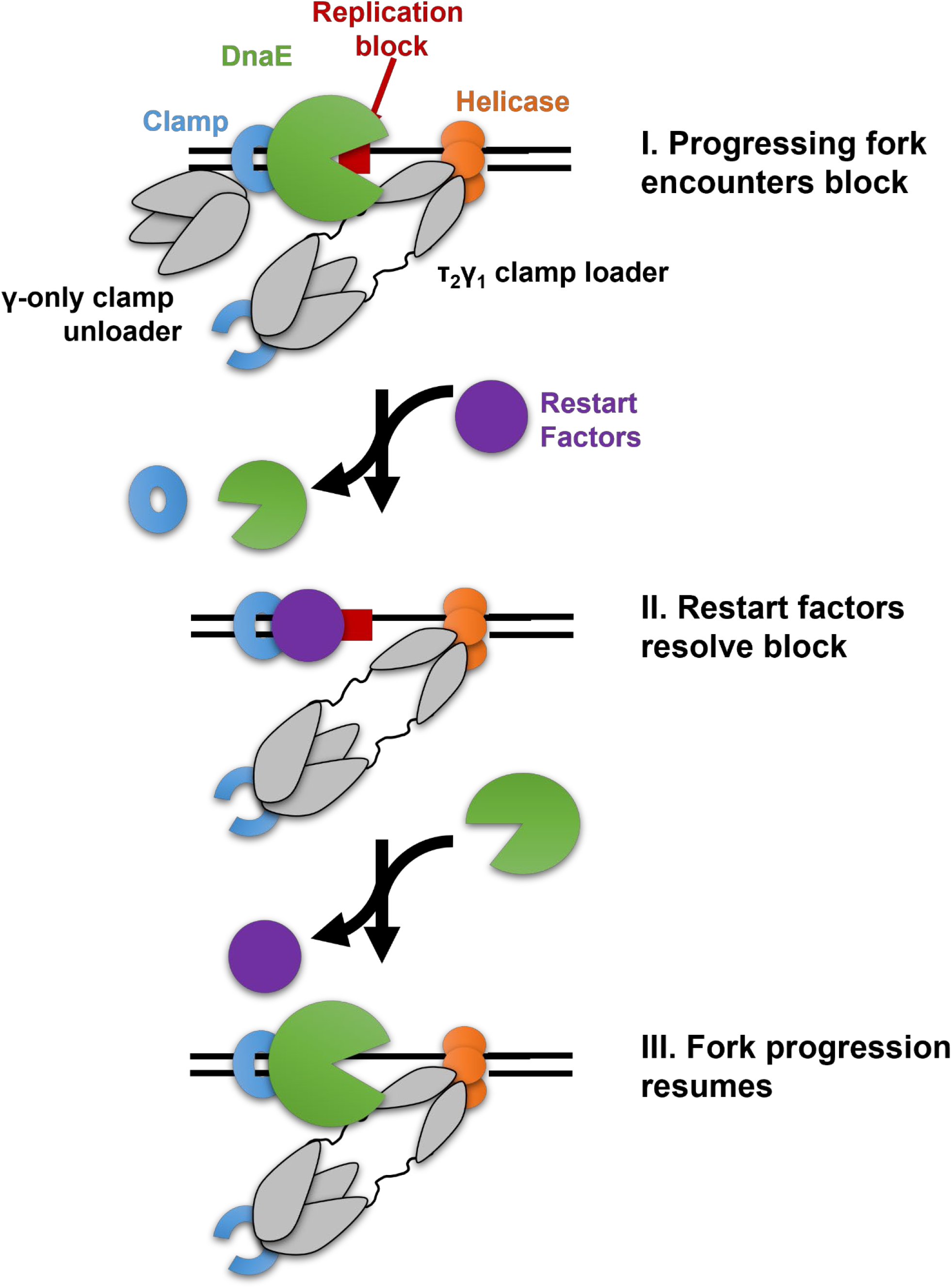
The bacterial clamp loader aids in resolution of replication fork stalling by allowing restart factors to access the replication fork. The DNA replication fork can encounter several types of replication blocks that cause replication fork stalling (e.g. DNA damage, collisions with transcription machinery, or physical tension caused by unwinding DNA). DnaX γ plays a role in allowing replication restart factors to access the replication fork (e.g. factors for DNA damage bypass (4), DNA repair (5–7), and DNA replication termination (8, 9)). This is critical for DNA replication restart and therefore cell survival.

The bacterial clamp loader is composed of five major subunits, three of which are DnaX (2, 11, 12). The single gene *dnaX* encodes two different forms of this subunit: the full-length τ and truncated γ. These forms were first discovered in *Escherichia coli*, where the γ form is created upon a ribosomal frameshift, which results in premature translation termination (13–15). While the two DnaX forms are conserved across many bacterial species, different bacteria create the γ form in different ways. For example, *Thermus thermophilus* creates a γ form upon RNA polymerase slippage, which results in premature transcriptional termination (16). Here we study *dnaX* in the alphaproteobacteria *Caulobacter crescentus*, where a full-length τ is translated and partially proteolyzed by the AAA+ protease ClpXP to create γ forms that lack the C-terminus (17).

The conservation of the γ form across bacteria suggests that is important to cell survival, however very little is known about its role. The DNA clamp loader complex binds the alpha subunits (DnaE) (18, 19) and helicase (DnaB) (20) of the DNA Polymerase III holoenzyme to coordinate replication on the leading and lagging strands (1). Since the C-terminal domain of DnaX is responsible for binding the DnaE (18), only the full-length τ can perform this job. Thus, out of the three DnaX subunits in the clamp loader complex, at least two of these must be τ to maintain two polymerases for lagging and leading strand synthesis. There is some debate in the field of whether this replicative clamp loader contains τ_3_ or τ_2_γ (4, 21–23).

While the γ subunit is not essential in *E. coli* (24), strains lacking γ are more sensitive to DNA damage and show a lower rate of DNA damage-induced mutagenesis (4). From this previous work it was suggested that *E. coli* γ plays a role in the DNA damage response, possibly by allowing the recruitment of error-prone DNA polymerases ((4), Figure 1). It has also been hypothesized that a γ-only clamp loader might be responsible for unloading clamps (25), which would be also useful in resolving stalled replication forks by removing the DnaE-bound DNA clamp (Figure 1). The elimination of γ from *E. coli* does not result in any observable phenotype in the absence of DNA damage (24). In contrast, our original characterization of *Caulobacter crescentus* suggested that γ was important even during normal growth (17).

Here, we investigate the consequences of changes in DnaX forms in *Caulobacter crescentus*. We demonstrate that switching from a processible form of DnaX to a non-processible variant (*τ-only* DnaX) is disfavored, but the strain grows normally once made. However, this strain is less tolerant of replication stress in the presence of DNA damaging agents. To explore this phenomenon further, we performed transposon sequencing to identify synthetic interactions with the *τ-only* allele, one of which was interruption of the damage-inducible cell division inhibitor *sidA*. Subsequent experiments show that the Δ*sidA τ-only* strain has a fitness defect compared to the wild type strain or single mutants under normal growth conditions and intolerance to DNA damaging agents. Consistent with this, we find that the *ΔsidA τ-only* strain has an elevated level of RecA protein and is dependent on *recA* for survival. This strain also has excess replication clamp foci during normal growth, suggesting a role for DnaX in preventing or resolving stalled replication forks, even in the absence of exogenous DNA damage treatment.

These data indicate that there is bias to maintain DnaX processing even in the absence of exogenous DNA damage. While *Caulobacter* can adapt to and tolerate the absence of DnaX processing, these cells are far less robust during DNA replication stress, during DNA damage treatment, or when DNA replication-cell division coordination is altered by deletion of *sidA*.

## MATERIALS AND METHODS

### Bacterial strains and growth conditions

Bacterial strains and plasmids used in this study are listed in Supplemental Table 1. All bacterial strains in this study are *Caulobacter crescentus* NA1000 derivatives and are grown in PYE medium (2 g/L peptone, 1 g/L yeast extract, 1 mM MgSO_4_, and 0.5 mM CaCl_2_) at 30°C. Solid media was made with 1.5% agar. The following antibiotics were used as indicated below: kanamycin (liquid culture – 10 μg/mL, solid media – 25 μg/mL), spectinomycin (liquid culture – 25 μg/mL, solid media – 200 μg/mL), oxytetracycline (liquid culture – 1 μg/mL, solid media – 2 μg/mL). Antibiotics were used when growing all strains containing self-replicating plasmids or single-recombination plasmid integrations.

### Constructing the *τ-only* strain using two-step recombination

The HinDIII-EcoRI fragment of pNPTS138 was replaced by a 2000 bp homologous region of the *Caulobacter* chromosome surrounding codons 544 and 545 of the *dnaX* coding sequence. These two codons were mutated by site directed mutagenesis (codon 544: GCG to GAC, codon 545: GCC to GAC) to replace DnaX alanine residues 544 and 545 with aspartic acid. This suicide vector was introduced into *Caulobacter* NA1000 by electroporation and primary integrations were selected for by plating on kanamycin. Single colonies were used to inoculate liquid cultures and lysates were analyzed by western blot to determine which form of DnaX was expressed. These liquid cultures were also plated on 3% sucrose to select for cells that had undergone secondary recombination to remove the plasmid sequence from the chromosome. Single isolates were then patched on antibiotic-free plates and on kanamycin to confirm the loss of the plasmid sequence. Kanamycin sensitive isolates were grown in liquid culture and lysates were analyzed by western blot to determine which DnaX form was expressed.

### Western blot

Western blots were performed using either an affinity purified rabbit *anti-Caulobacter* DnaX primary antibody (1:10,000 dilution, 1 hour) or a commercially available rabbit anti-*E. coli* RecA primary antibody (1:10,000 dilution; Abcam, Cambridge, MA, 16 hours). Blots were visualized using the goat anti-rabbit Alexa Fluor^™^ Plus 800 secondary antibody (1:10,000 dilution; Licor, Lincoln, NE, 1 hour). For normalization purposes, blots were then probed with a rabbit anti-*Caulobacter* ClpP serum (1:10,000 dilution, 16 hours) and visualized with the same Licor secondary antibody. Bands were quantified using ImageJ (26).

### Bacterial growth curves

Cells in stationary (Figures 3B and 7B) or exponential phase (Figures 4B and 5B) were diluted to an OD600 = 0.1 in a 96-well plate. Cells were grown shaking at 30°C in a platereader and OD600 was measured every 20 minutes for 15 hours. To find growth rate, the slope of the most linear portion of each replicate was taken.

**Figure 2.**
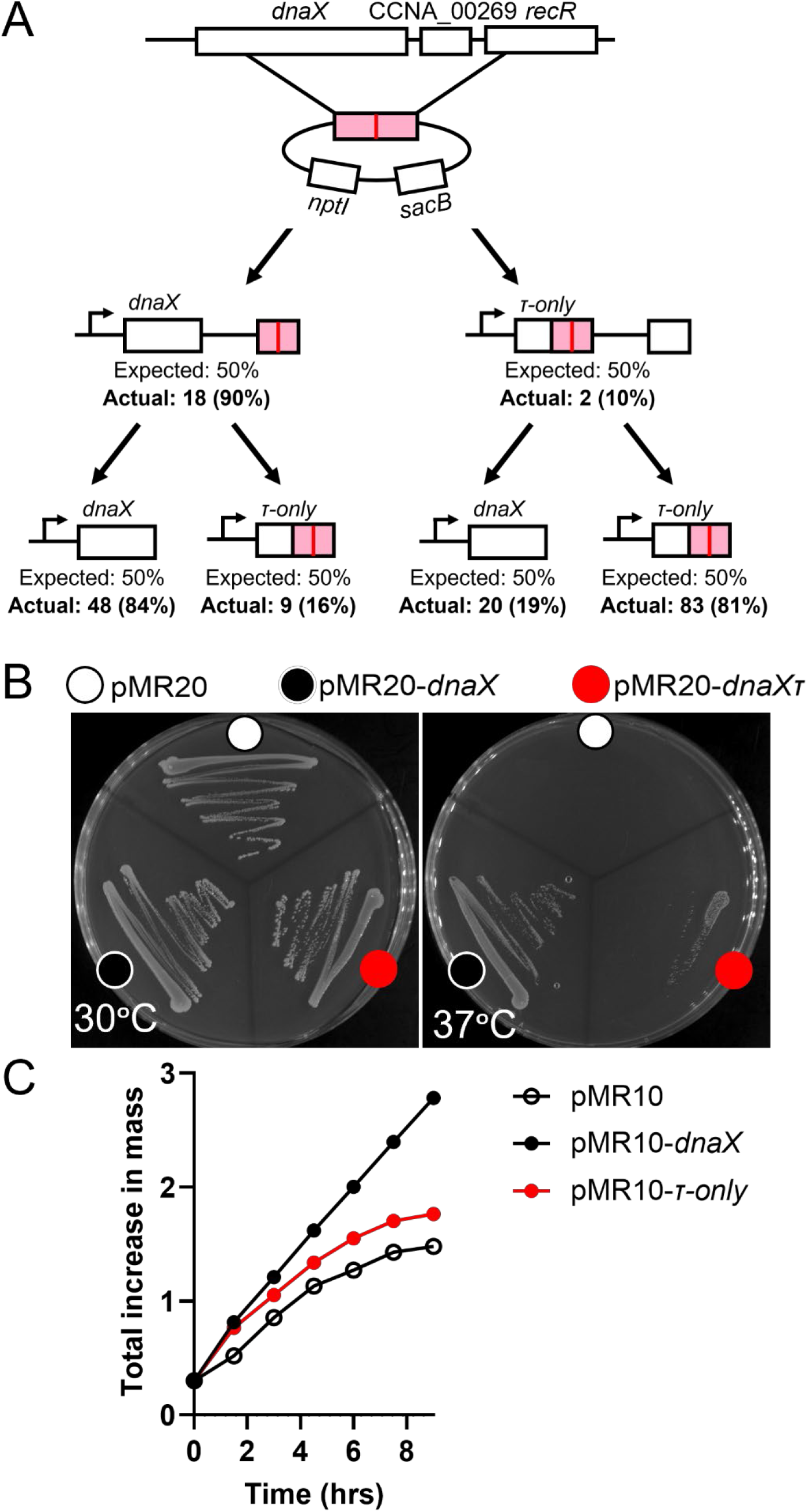
There is a bias against eliminating DnaX processing, but rare isolates make strain construction possible. (A) We used a two-step recombination strategy to construct the *τ-only* strain. The suicide vector was designed to produce a 1:1 ratio of wild type *dnaX* to *τ-only* allele expression at the primary integration and secondary recombination steps, however both steps favored maintaining the wild type *dnaX* over recombining to switch to expression of the *τ-only* allele. (B) While the wild type *dnaX* allele rescues growth of the *dnaX_τs_* strain at nonpermissive temperature, the *τ-only* allele was not able to fully rescue. Experiment performed in triplicate, representative images shown. (C) Overnight growth was back diluted to OD600=0.3 and grown at non-permissive temperature for 1.5 hours. OD600 was measured and cells were back diluted to OD600=0.3. Measurement and back dilution were repeated every 1.5 hours for 9 hours. A cumulative change in OD600 was calculated at each timepoint. Results of a single experiment are shown. As in the plating assay, the wild type *dnaX* allele rescued the growth of the *dnaX_Δs_* strain at nonpermissive temperature, but the *τ-only* allele did not.

**Figure 3.**
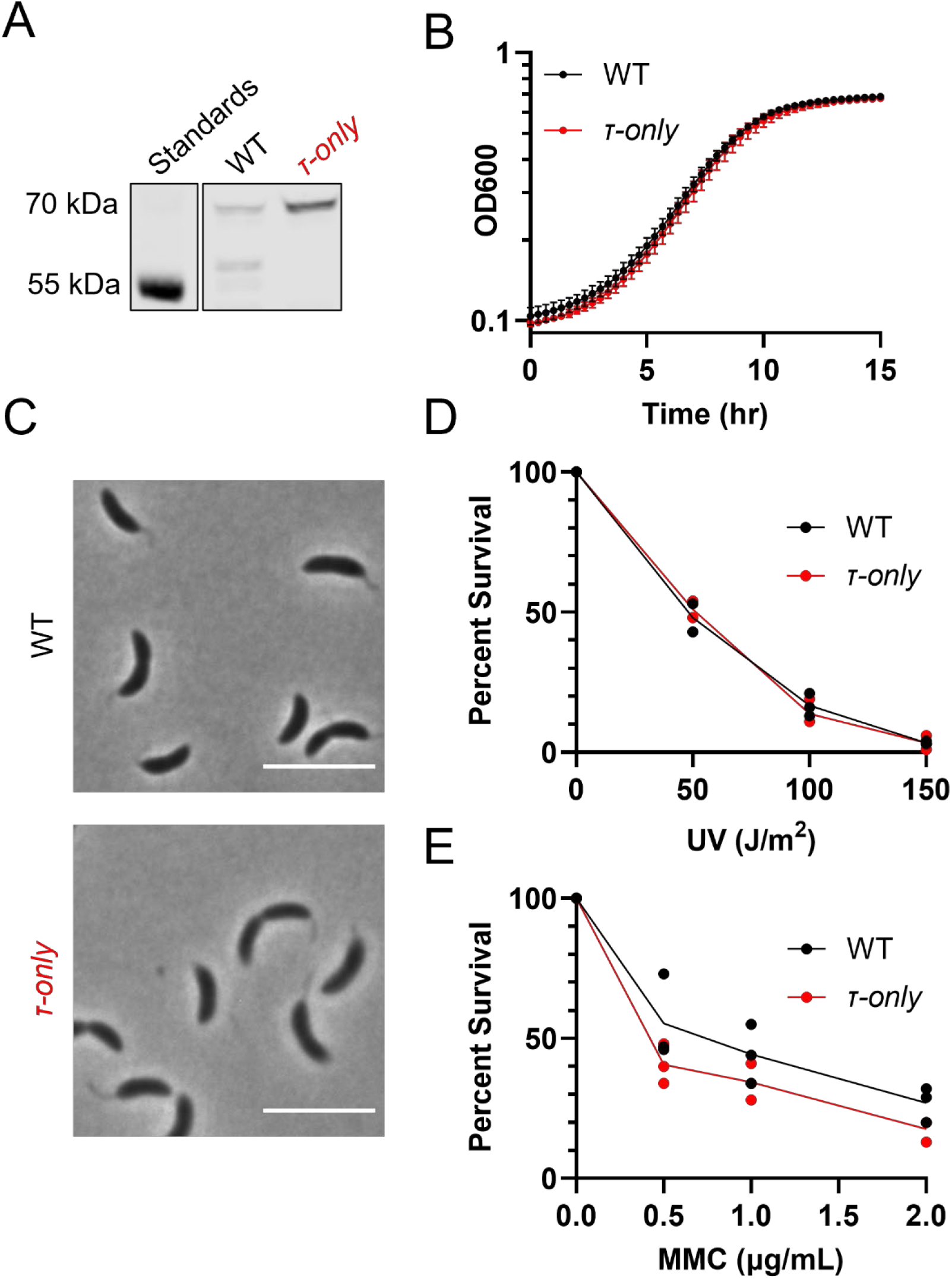
Absence of DnaX processing does not affect cell growth or morphology. (A) The final *τ-only* strain was analyzed by western blot using the *Caulobacter* anti-DnaX antibody. Instead of the three DnaX forms observed in the wild type strain, only the longest DnaX form (τ) is observed. Growth (B) and morphology (C) of this strain is comparable to the wild type strain. Experiments in (B) performed in triplicate, mean and standard deviation are shown. Where error bars are smaller than the width of the marker, error bars were omitted. Experiment in (C) was performed in triplicate, representative images shown. Scale bars represents 5μm. (D and E) Exponential phase cells were treated with various amounts of UV light (D) or varying concentrations of mitomycin C for one hour (E) The τ-only strain demonstrated a consistent, but mild defect in mitomycin C survival, but not in survival to ultraviolet light. Experiments performed in triplicate, mean and individual data shown.

**Figure 4.**
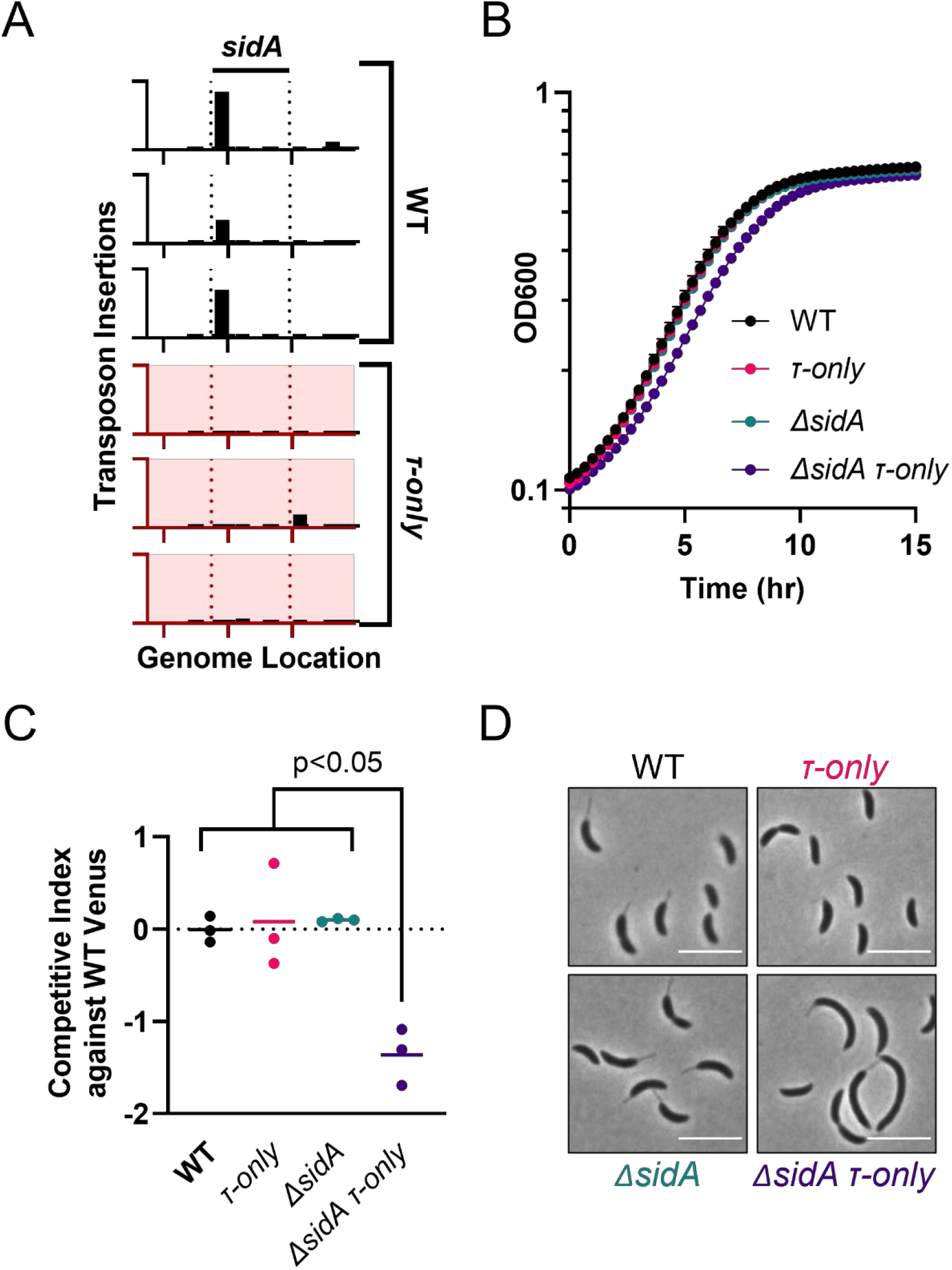
Cell division inhibitor *sidA* is important in the absence of DnaX processing. (A) A transposon-sequencing experiment showed fewer insertions in the gene *sidA* in a *τ-only* background compared to in a wild type background. The y-axis of each plot measures the number of transposon insertion on a linear scale from 0-200. Each x-axis shows genome location of nucleotides 2152211-2152696 in *Caulobacter* strain NA1000 **(40)**. Deletion of *sidA* in a *τ-only* strain results in a growth defect (B and C) and filamentation (D). All experiments were performed in triplicate. In (B) mean and standard deviation are shown. Where error bars are smaller than the width of the marker, error bars were omitted. Growth rate quantification is shown in Supplemental Figure 2A. The *ΔsidA τ-only* strain has a significantly lower slope than each of the other strains (p-values <0.05). (C) When each test strain is competed against in co-culture a wild type Venus-expressing fluorescent reporter, only the *ΔsidA τ-only* mutant shows a competitive disadvantage. Mean and results of two-tailed t-test are shown. This is consistent with the growth defect observed in B. In (D) representative images are shown. Quantifications of cell length are shown in Supplemental Figure 2B. Scale bars represent 5μm.

**Figure 5.**
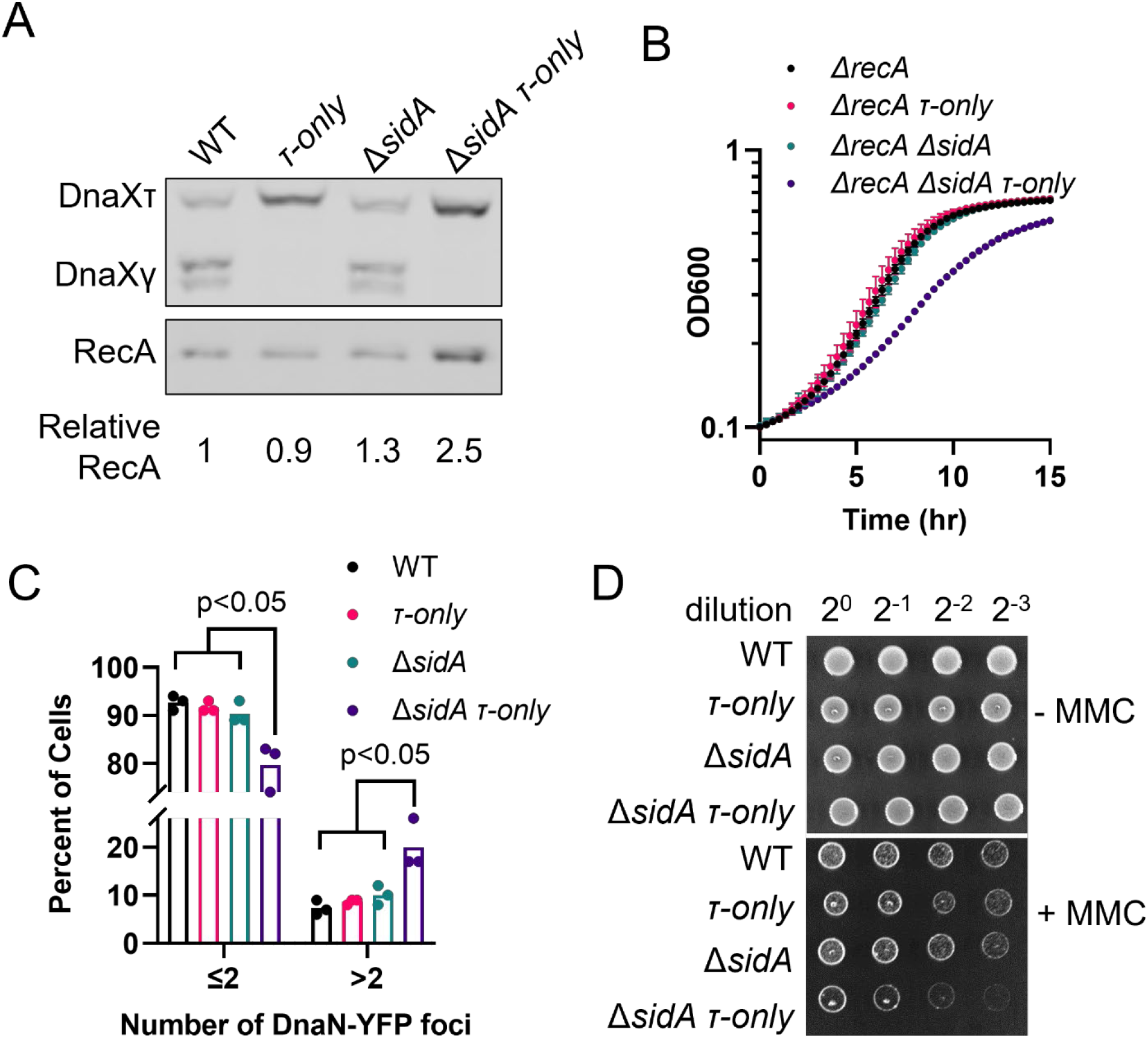
The *τ-only* allele and *sidA* deletion results in a synthetic phenotype of a deficiency in resolving replication fork stalling events. (A) Western blot analysis shows an increased level of RecA protein in the *ΔsidA τ-only* strain compared to the wild type or single mutant strains. Variability of relative RecA concentration is approximately 10%. Full quantification of RecA levels shown in Supplemental Figure 3A. A comparison of a growth curve of these parental (*recA*+) strains (Figure 4B) and *recA* knockouts in these strains (B) indicates that *recA* is critical for growth of the *ΔsidA τ-only* strain compared to the wild type or single mutant strains. Mean and standard deviation are shown. Where error bars are smaller than the width of the marker, error bars were omitted. Quantification of these growth rates shown in Supplemental Figure 3B (C) When DnaN-YFP foci are observe in the parental (*recA*+) strains, the *ΔsidA τ-only* strain has an increased proportion of cells that contain >2 DnaN-YFP foci. This indicates a greater than normal number of replisomes. The extra foci are likely due to replication-dependent DNA repair. All experiments were performed in triplicate. Mean and standard deviation shown. (D) We find that the *ΔsidA τ-only* is more sensitive to mitomycin C than the wild type strain of single mutants. Cells were spotted on 0 or 1 μg/mL mitomycin C. Spots in a row represent 2-fold serial dilutions. Representative images of triplicate experiments shown.

**Figure 6.**
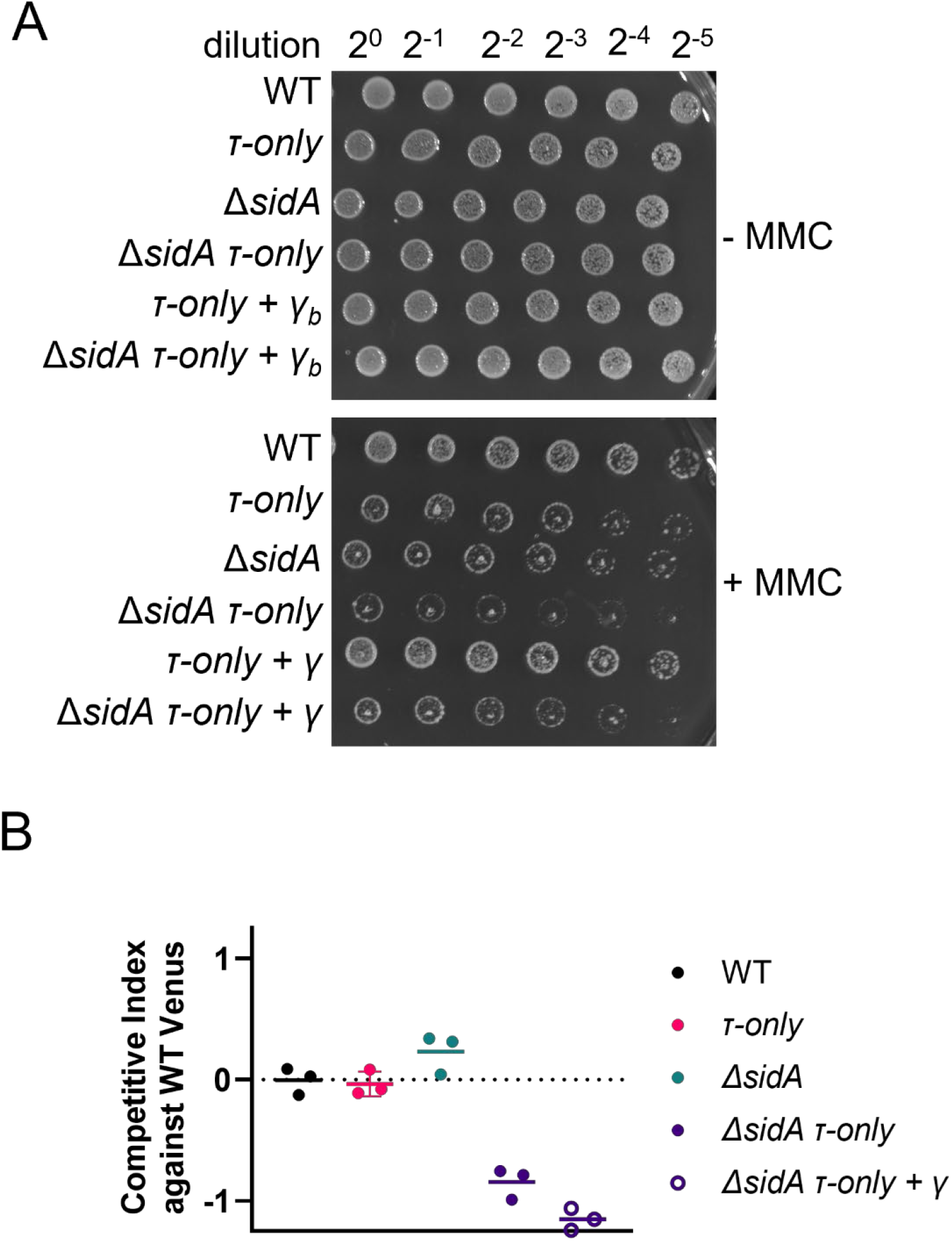
Mitomycin C sensitivity in the *ΔsidA τ-only strain is partially complemented by expression of* γ, but poor competition is not. (A) The MMC sensitivity in the *ΔsidA τ-only* strain is partially complemented by expressing of γ (amino acids 1-492) at an alternate locus (xylose locus). Cells were spotted on 0 or 1 μg/mL mitomycin C. Spots in a row represent 2-fold serial dilutions. Representative images of triplicate experiments shown. Other replicates shown in Supplemental Figure 5. (B) The competitive disadvantage of the *ΔsidA τ-only* strain is not complemented by expression γ (amino acids 1-492) at an alternate locus (xylose locus).

**Figure 7.**
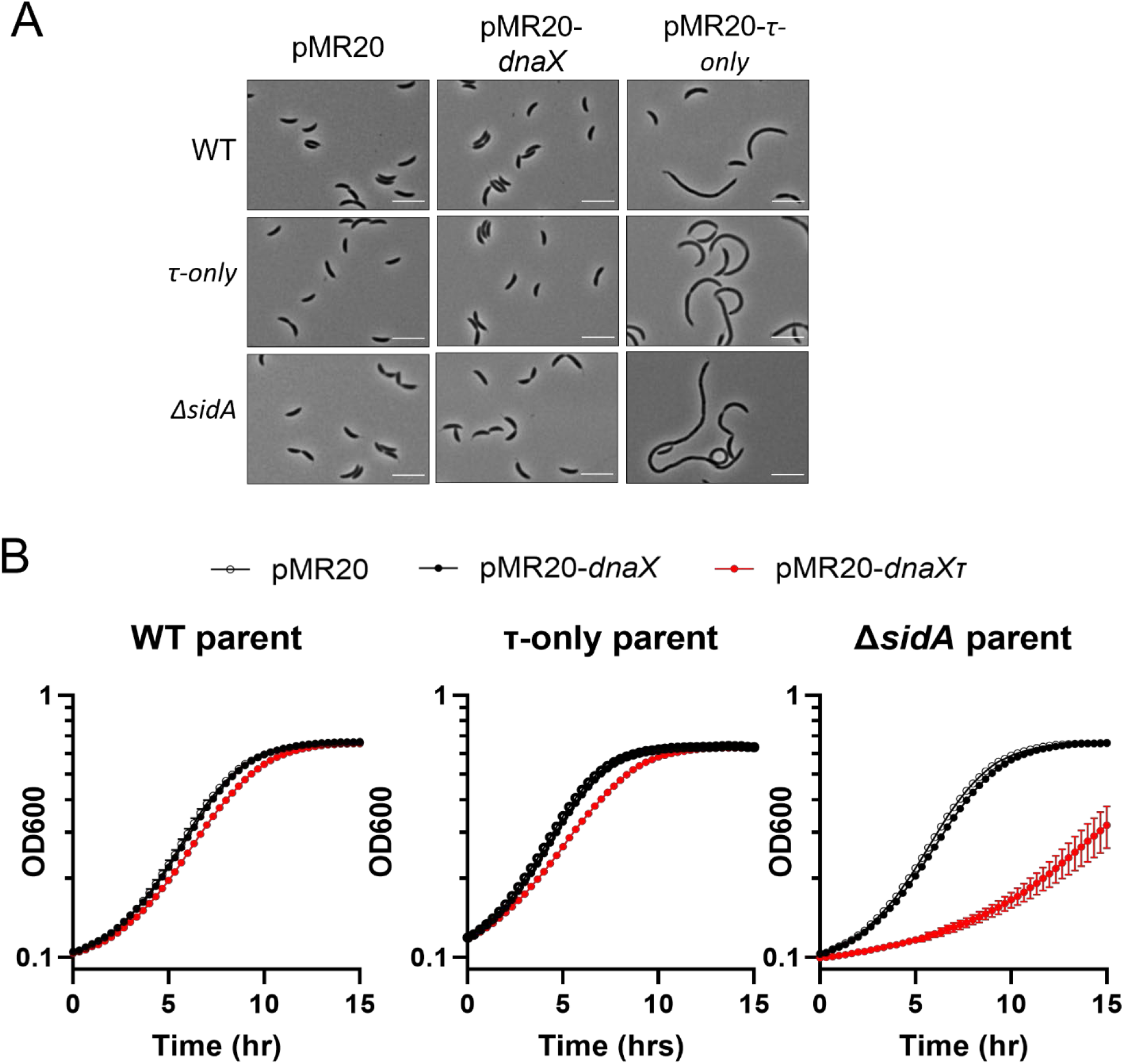
Excess non-processible DnaX is toxic in the absence of *sidA*. Introduction of excess *τ-only* results in (A) elongation and (B) slower growth of *ΔsidA* cells, while introduction of excess wild type *dnaX* has no significant effect. All experiments performed in triplicate. In (A) representative images shown, quantification of cell length shown in Supplemental Figure 6A. In (B) mean and standard deviation are shown. Where error bars are smaller than the width of the marker, error bars were omitted. Growth rate quantification shown in Supplemental Figure 6B.

### Cell morphology measurements and competition assays

For cell morphology measurements, cells were back diluted and grown for 10-15 generations to an OD600 of about 0.5. Cells were imaged by phase contrast microscopy under 1000X magnification. ImageJ (26) and MicrobeJ (27) were used to identify cells and quantify cell length.

Overnight cultures of each test strain and the Venus-expressing reporter strain were mixed 1:1 according to OD600 and serially diluted 30,000-fold in a final volume of 10 mL PYE. Undiluted cells were imaged by phase microscopy and fluorescence microscopy using a GFP-selecting filter. ImageJ was used to manually quantify the number of fluorescent and non-fluorescent cells in the images for each sample. Diluted samples were grown to stationary phase and imaged again. The competitive index was calculated as the log_2_[((ratio test/Venus)_END_ / (ratio test/Venus))_START_ / ((ratio wild type/Venus)_END_/[(ratio wild type/Venus))_START_].

### DNA damage susceptibility testing

For the acute stress testing (Figure 3D and E), cells were grown to exponential phase and treated with UV light using a Stratalinker UV crosslinker or with mitomycin C (Millipore Sigma, St. Louis, MO) for one hour. Cells were serially diluted in PYE, plated on PYE agar, and incubated at 30°C for 48 hours. Colony forming units were counted and the CFU count of the original sample was calculated as (number of CFU on plate)*(dilution factor). Percent survival was calculated as CFU count for treated cells / CFU count for untreated cells.

For chronic stress (Figures 5D and Supplemental Figure 5), stationary phase cultures were normalized to an OD600 = 0.5, diluted 100-fold (left-most spot in a row) and then serially diluted by 2-fold for 7 dilutions, and plated on PYE agar and PYE agar + mitomycin C. Plates were incubated for 48 hours (untreated) or 60 hours (treated) at 30°C, then imaged using a Syngene G:Box.

### Temperature sensitivity complementation

For plate-based method, colonies from fresh plates were struck onto PYE agar and inoculated at permissive temperature (30°C) or nonpermissive temperature (42°C).

For liquid-based method, overnight growth was diluted and grown to exponential phase at permissive temperature. At time 0, cells were normalized to an OD600 = 0.3 and shifted to nonpermissive temperature. Every 2 hours, OD600 was measured, and cells were diluted back to OD600 = 0.3 in pre-warmed PYE. The total increase in mass was calculated as the cumulative increase in OD600.

### Transposon sequencing

Transposon libraries were created using the wild type and *τ-only* parental strains as previously described (28, 29). The wild type and *τ-only* libraries outgrown and sequenced in triplicate. Data was analyzed using edgeR(30). Differential analysis of the counts in these libraries are available in Supplemental Table 2. The gene *sidA* (CCNA_02004) showed a significantly lower number of insertions in the *τ-only* library compared to the wild type library (*log2*(*τ-only/wild type*) = ~4.8, See Figure 4).

### Quantifying DnaN-YFP foci

We used the single integration plasmid pNABC198 (a generous gift from the Badrinarayanan lab; 10) to tag DnaN with a C-terminal YFP tag at the native locus in the wild type, *τ-only, ΔsidA*, and *ΔsidA τ-only* parental strains. Cells were grown to exponential phase and imaged with phase contrast and fluorescence microscopy. YFP foci were quantified manually using ImageJ (26).

## RESULTS

### A bias against the τ-only allele revealed during strain construction

Our lab has previously reported on a form of DnaX that is not proteolyzed by ClpXP (DnaXnp, 8). This DnaX variant contains two frameshift mutations that alters the amino acid sequence of a large portion of DnaX adjacent to the predicted ClpXP-recognition site (17). This variant is recognized by ClpXP but fails to generate partial proteolysis fragments. To completely eliminate ClpXP recognition we substituted two adjacent amino acids (AA544 and 545 to DD) in the predicted ClpXP recognition site, with the rest of the sequence remaining intact. This variant is unable to be processed at all by ClpXP in vitro (Supplemental Figure 1).

A two-step double recombination method was used to replace the *Caulobacter dnaX* gene with the *τ-only* allele. We first engineered point mutations in the center of a 2000bp region of homology cloned into the pNPTS138 suicide vector which cannot propagate in Caulobacter and contains a counter-selectable *sacB* marker along with a kanamycin resistance cassette ((3), Figure 2A, top). The homology region starts at nucleotide 630 in the *dnaX* coding sequence (1000 bp upstream of the mutations) and ends 822 nucleotides after the end of the *dnaX* reading frame (1000 bp downstream of the mutations). Based on sequence alone, approximately half of primary integrations were expected to occur within the homology region before the mutations, leading to expression of the *τ-only* allele from the chromosomal promoter (Figure 2A, middle). The other half were expected to occur in the homology region after the mutations, resulting in expression of the wild type *dnaX* allele from the chromosomal promoter (Figure 2A, middle). Western blot analysis was used to determine the percentage of isolates expressing wild type *dnaX* and the percentage expressing *τ-only*. The wild type *dnaX* allele expresses three DnaX forms, while the *τ-only* allele expresses only the largest DnaX form (Figure 3A). Surprisingly, about 90% of primary integration isolates (18/20) expressed the wild type *dnaX* allele from the chromosomal promoter (Figure 2A, middle). Since there is no reason to believe that a bias exists in the locations of recombination events, this suggests a strong bias exists against eliminating DnaX processing.

Secondary recombination events that had eliminated the plasmid from the chromosome were then identified by growing cells in nonselective media and plating on sucrose containing media, which selects for loss of *sacB* function, and then verifying kanamycin sensitivity, which indicated loss of the *nptI* gene function. Because *dnaX* is essential, these isolates must have kept one of the two homologous regions surrounding the *dnaX* mutation site but lost the plasmid sequence with the second homologous sequence. Based on a model where the secondary recombination occurs randomly, the secondary recombinants were expected to have a roughly equal probability of maintaining either the *dnaX* or *τ-only* allele, despite which allele was expressed in the primary integration (Figure 2A, bottom). When a *dnaX*-expressing primary integrant was used, almost 90% of isolates expressed wild type *dnaX* (48/57; Figure 2A, bottom left). One of the *τ-only-* expressing isolates from this process is used in the remainder of this report and is called the *τ-only* strain.

When a *τ-only*-expressing primary integration was used for secondary recombination, there was a strong bias for maintaining the *τ-only* allele (83/103; Figure 2A, bottom right). Five of the final *τ-only* strains were analyzed using short-read whole genome sequencing, which did not reveal any suppressor mutations (See Supplemental Tables 3-7). These data suggest that eliminating DnaX processing by converting from a wild type *dnaX* to the *τ-only* allele is a rare event, but that *Caulobacter* can adapt to expression of the *τ-only* allele without the presence of a suppressor mutation.

To explore this phenomenon further, we introduced plasmid-borne copies of these alleles into a temperature sensitive strain of *dnaX*. We had previously reported that the *dnaX_np_* allele was unable to rescue growth of a temperature sensitive allele of *dnaX* (17). It was unclear whether the lack of processing itself or the large change in amino acid sequence of the DnaX_np_ was the cause of this phenotype. Unlike the allelic replacement result, we find that the *τ-only* allele is insufficient to completely rescue growth of the *dnaX_TS_* strain at nonpermissive temperature (Figure 2B and C). These data support our observation above that the transition from wild type *dnaX* to the *τ-only* allele is detrimental, a consequence accentuated during acute loss of DnaX activity in a temperature sensitive background. These data also align with what had previously been observed by our lab for the *dnaX_np_* allele(17).

### The *τ-only* strain is mildly sensitive to DNA damage by mitomycin C, but not ultraviolet light

Despite the apparent bias observed in its construction, the *τ-only* strain shows no defects in growth or morphology (Figure 3B and C). *E. coli τ-only* strains are sensitive to ultra-violet (UV) radiation (4), but the *τ-only* strain of *Caulobacter* does not show the same sensitivity (Figure 3D). This difference might be explained by the fact that over *Caulobacter’s* evolution, it has naturally been exposed to high doses of ultraviolet light in its native freshwater environment. The *τ-only* strain does exhibit a mild sensitivity to mitomycin C, with approximately 65-75% the survival rate to mitomycin C treatment of the wild type cells (Figure 3E). This evidence suggests that, while the cells can adapt to the absence of DnaX processing, this change leaves them susceptible to certain forms of DNA replication stress, such as by treatment with mitomycin C.

### Deletion of cell division inhibitor *sidA* is detrimental in the absence of DnaX processing

To gain insight into the role of DnaX processing, we performed transposon sequencing to identify synthetic interactions that would help us determine how cells are adapting to the *τ-only* allele. We created and sequenced EZ-Tn5 transposon libraries in the wild type and *τ-only* backgrounds, then compared the distribution of insertions. Each library contained approximately 200,000 unique insertion sites ((29) and see Methods).

We found significantly smaller number of insertions in the *sidA* gene for the *τ-only* background compared to the wild type background (log2(fold difference) = −4.8). We found that the bulk of the *sidA* insertions in the wild type library were located at a single site (Figure 4A), which was not surprising due to the small size of the *sidA* coding sequence (123 bp). There were no insertions at this site in the *τ-only* library. The *sidA* gene encodes a LexA/RecA-controlled protein that is thought to inhibit cell division during the DNA damage response (33). SidA is one of two DNA damage-induced *Caulobacter* cell division inhibitors, which are thought to stall cell division and allow cells to repair damaged DNA before it is passed on to a daughter cell (33, 34).

We constructed a *ΔsidA τ-only* strain and found that this strain grows more slowly than either parent or wildtype (Figure 4B, Supplemental Figure 2A). To explore this growth defect further, we used a co-culture competition experiment. The *ΔsidA τ-only* strain has a significantly lower ability to compete with a fluorescently labelled wild type reporter strain than the wild type strain or single mutant strains (Figure 4C). To confirm that this is not an artifact of the wild type fluorescent reporter strain, we also tested the single mutants against a *ΔsidA τ-only* fluorescent reporter and vice versa (Supplemental Figure 2C and D). In both cases, the individual mutant strains have a higher ability to compete in co-culture than the *ΔsidA τ-only* strain (Supplemental Figure 2C and D).

### τ-only ΔsidA cells show increased RecA dependence

The *ΔsidA τ-only* cells are also longer (Figure 4D, Supplemental Figure 2B) than the single mutants and wild type strain. As cell filamentation can be a sign of cellular distress such as that seen during DNA damage (33), we hypothesized that deletion of the *sidA* cell division inhibitor affects coordination between DNA replication and cell division, leading to increased DNA replication stress in the absence of DnaX processing. Consistent with this interpretation, RecA levels in the double *ΔsidA τ-only* mutant were elevated (Figure 5A, quantification in Supplemental Figure 3A). Furthermore, deletion of *recA* in this strain severely inhibited growth (Figure 5B, Supplemental Figure 3B), indicating that a functional DNA damage response is required for normal growth of the double *ΔsidA τ-only* mutant. These data suggest that the cell division inhibitor *sidA* is necessary in the absence of DnaX processing likely due to DNA replication stress. SidA and its function as a cell division inhibitor are likely one mechanism by which the cell adapts to the *τ-only* allele during this stressful time.

To directly obtain insight into replisome dynamics in the *ΔsidA τ-only* strain, we monitored fluorescently labelled DNA clamp (DnaN-YFP) foci as a proxy for replisomes. While lone DNA clamps slide along DNA of diffuse in solution freely, replisome-associated clamps move slowly enough to form fluorescent foci. In asynchronous wild type and single mutant strains, most cells had 1-2 replication foci likely corresponding to active replication forks and/or a small number of DNA damage-related foci (e.g. stalled replication forks, lesion bypass or lesion repair sites). A small population of cells contain more than two DnaN foci (<10%, Figure 5C). These extra foci are likely to be related to DNA damage or replication fork stalling, since only a single DNA replication initiation event occurs in one *Caulobacter* cell cycle. Importantly, we find that the *ΔsidA τ-only* strain has twice as many cells with >2 foci than the other strains (~20%; Figure 5C). Given the RecA induction and importance of *recA* to the growth of this strain, these extra foci are likely related to DNA damage and/or replication fork stalling. Consistent with this interpretation, the strain is more sensitive to mitomycin C than the wild type strain or single mutants (Figure 5D).

These data can be explained by two possibilities: either the *ΔsidA τ-only* strain has a higher amount of damaged DNA, even in the absence of an exogenous DNA damaging treatment, or DNA damage-related foci are more persistent in this strain. Either possibility explains the filamentation, RecA induction, and dependance on *recA* that we observed above. While *Caulobacter* can tolerate either the absence of DnaX processing or the absence of the cell division inhibitor *sidA*, both mutations lead to an overwhelming number of DNA damage or replication fork stalling events.

### Phenotypic defects arise from both lack of γ and surplus of τ

Since the *τ-only* strain has a higher total amount of DnaX in the cell (Supplemental Figure 4), it is unclear whether bias against the *τ-only* allele and the phenotypes observed in the *τ-only* and *ΔsidA τ-only* strains are due to the absence of the γ forms or due to the increased abundance of DnaX overall in the strain. To address this question, we attempted to complement the *ΔsidA τ-only* strain’s mitomycin C sensitivity by expressing a truncated DnaX form that mimicked the estimated size of the longer *Caulobacter* γ form. The MMC sensitivity of the *ΔsidA τ-only* strain is partially rescued by expression of γ (DnaX(1-492)) at a second locus (Figure 6A).

Next, we tried to complement the *ΔsidA τ-only* strain’s inability to compete in coculture by expressing γ (DnaX(1-492)), but γ expression did not complement this phenotype (Figure 6B). These data indicate that while the absence of γ is likely involved in the phenotypes of the *ΔsidA τ-only* double mutant, it is not the full explanation for these phenotypes.

Our results so far suggests that a surplus of the *τ*-form of DnaX may be toxic to cells lacking *sidA*. To explicitly test this, we introduced a second copy of *dnaX* on a low copy plasmid into the wild type, *τ-only*, and *ΔsidA* strains. In all parental strains, addition of an exogenous *τ-only* allele resulted in a significant increase in cell length (Figure 7A and Supplemental Figure 6A) and decrease in growth rate (Figure 7B, Supplemental Figure 6B, and Supplemental Table 8), while an exogenous copy of the wild type *dnaX* had little to no effect. Additionally, we find that the effects of an exogenous *τ-only* allele on cell length and growth rate are much larger in the *ΔsidA* strain than in the wild type and *τ-only* strains (Figure 7A and B, Supplemental Figure 6A and B, and Supplemental Table 8), despite total DnaX levels being similar in the two parental strains for each plasmid (Supplemental Table 5C).

These data suggest that phenotypes associated with the *τ-only* allele are not simply due to the absence of γ. They also suggest that the bias shifting to the *τ-only* allele in strain construction was not due to a simple elevation in total DnaX levels. It appears that an excess of a non-processible form of DnaX is detrimental to the cell, particularly in the absence of *sidA*. We propose that DnaX processing is important to preventing or resolving instances of replication fork stalling and that *sidA* can compensate for the loss of DnaX processing, likely by adjusting cell division. An excess of *τ-only* DnaX interferes with this role and results in a sensitivity to DNA replication stress.

## DISCUSSION

Here, we find that there is selection to maintain DnaX processing in *Caulobacter*. Though *Caulobacter* can survive without this processing, they are less able to survive stress related to DNA replication. These findings are in line with previous work in *E. coli*, where ribosomal frameshifting during DnaX translation aids survival of exogenous sources of DNA damage (4). This need for γ is explained by the higher preference of *E.coli τ-only* clamp loader for DnaE, the replicative polymerase, which restricts access for alternative DNA polymerase loading during replication fork stalling (4). Our results here show increased DnaN-YFP foci, dependence on recA, and increased mitomycin C sensitivity in the *ΔsidA τ-only* strain suggests that the *Caulobacter τ-only* clamp loader also has difficulty in resolving replication fork stalling events and that the clamp loader likely plays a similar role in *Caulobacter* as it does in *E. coli* (Figure 8).

**Figure 8.**
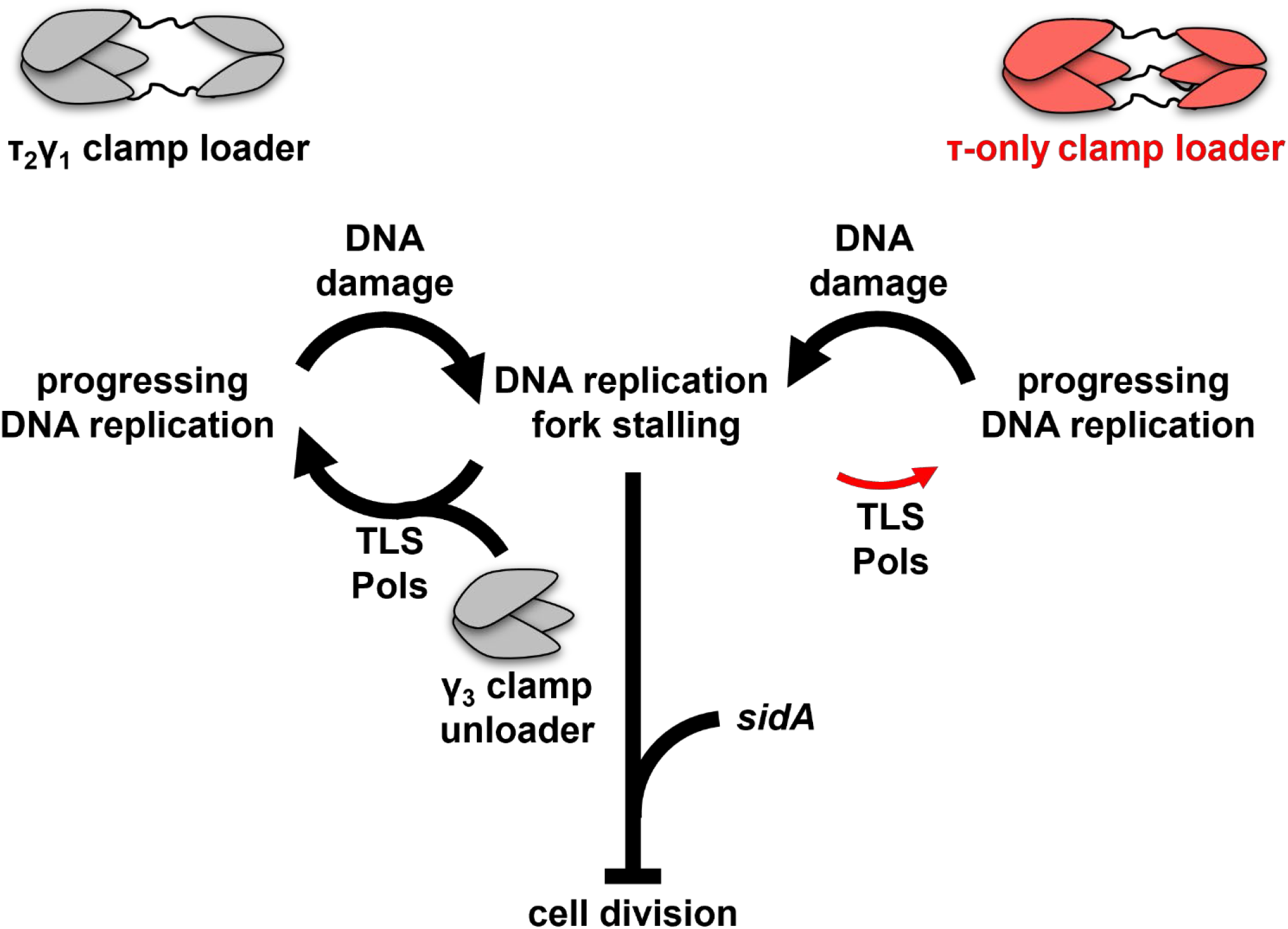
Clamp Loader composition affects resolution of stalled replication forks. Our data supports the model that the τ-only clamp loader is less effective at allowing alternative translesion (TLS) DNA polymerases and possibly other replication restart factors (e.g. DNA repair factors, DNA replication termination factors) to access the fork than the wild type τ2γ1 clamp loader. This deficient access of replication restart factors causes a build-up of stalled replication forks, which triggers a *sidA*-dependent mechanism to compensate for this, likely involving inhibition of cell division. The expression of γ in the τ-only strain improves its DNA damage tolerance, possibly due to a γ-only clamp unloader assisting in removal of stalled DnaE-associated DNA clamps, which allows some restart machinery to better access the clamp.

We find that expression of γ can partially rescue the mitomycin C sensitivity in the *ΔsidA τ-only* but cannot suppress this strain’s reduced fitness under unstressed conditions. It has been suggested that γ could primarily participate in clamp unloading (25) and eukaryotes encode the clamp unloader Elg1/ATAD5 that removes PCNA clamp at stalled replication sites during DNA damage (reviewed in (35)). Based on this, we speculate that a γ-only clamp unloader is providing some relief to the mitomycin C-treated *ΔsidA τ-only* strain by helping to unload clamp and DnaE at stalled replication forks. This would allow replication restart machinery to access and repair or bypass the DNA damage.

By contrast, expression of γ does not rescue the *ΔsidA τ-only* failure to compete with wildtype in the absence of DNA damage. Our interpretation that overabundance of *τ-only* DnaX or an imbalance of τ to γ ratio may play a role in this phenotype. We note that excessive *τ-only* results in slower growth even in wildtype cells (Figure 7B), suggesting that an excess of the DnaX C-terminal domain may have negative effects even in wildtype backgrounds, such as titration of the DnaE away from the replication fork.

The mitomycin C sensitivity of the *τ-only* strain is amplified when *sidA* is deleted, with defects in growth and dependence on *recA* that are not seen in the parent strains. While *sidA* is highly upregulated during DNA damage (33), cells lacking *sidA* show no deficiency in DNA damage tolerance (Figure 5D), presumably due to redundant pathways such as *didA* (34). The only published effect for *sidA* alone is that overexpression is sufficient to halt cell division, even in the absence of DNA damage (33). We hypothesize that *sidA* inhibition of cell division is required in the *τ-only* strain to compensate for deficiencies in clamp loader activities such as alterative polymerase switching, needed for repair under DNA damaging conditions (Figure 3E, Figure 6; Figure 8). Similarly, elevating *τ* levels in a *ΔsidA* results in a severe growth defect and filamentation (Figure 7), consistent with a need for *sidA* to adjust cell division in response to DNA replication needs.

Our overall model is that the *τ-only* clamp loader is not as effective in its role in replication restart. This may be because the *τ-only* clamp loader is less effective in allowing restart factors, such as alternative DNA polymerases, to access the fork (Figure 8), as has been found for the *τ-only* clamp loader and alternative DNA polymerases in *E. coli* (4). The switch of wild type clamp loaders to *τ-only* clamp loaders is initially very detrimental, as we see in our strain construction (Figure 2A) and temperature sensitivity complementation (Figure 2B and C). Though there is a bias against switching to the *τ-only* allele, cells can adapt and grow normally (Figure 3B and C). One such adaptation seems to rely on *sidA* to adjust the coordination of DNA replication and cell division (Figure 7).

While our results indicate that the capacity to process DnaX is important, particularly during DNA replication stress, it also shows that DnaX processing is not essential for cell survival and that cells can eventually adapt to a *τ-only* allele. This may explain why the τ and γ forms of DnaX are generally conserved in bacteria, but that the pathways to the shorter γ form varies. A possible scenario is that early bacteria used a clamp loader with identical DnaX subunits, which would have more closely resembled the T4 phage clamp loader (reviewed in (36)). Over time, perhaps coinciding with the increased risk to DNA damage that came with an oxygen-rich atmosphere, bacteria may have independently evolved mechanisms to create γ. Eukaryotic clamp loaders took a different route to evolve a similar system, where the multiple DnaX-like subunits of the clamp loader are encoded by unique genes. This might also explain why there are bacterial species that do not appear to produce a γ form, such as *Streptococcus pyogenes, Aquifex aeolicus*, and *Bacillus subtilis* (37–39).

## ACKNOWLEDGEMENTS

This work was supported by NIH/NIGMS R35GM130320 to P. Chien. The *pdnaN-yfp* integrating plasmid was a generous gift from the Badrinarayanan Lab at the National Centre for Biological Sciences – Tata Institute of Fundamental Research, Bangalore, India. The ΔsidA strain was a kind gift from the Laub lab at the Massachusetts Institute of Technology. The authors would like to thank the members of the Chien lab and the Goley lab at Johns Hopkins University, Baltimore, MD, USA for helpful discussions about this work.

## Notes

### Competing Interest Statement

The authors have declared no competing interest.

