## Supplemental Figures for "Clamp loader processing is important during DNA replication stress"

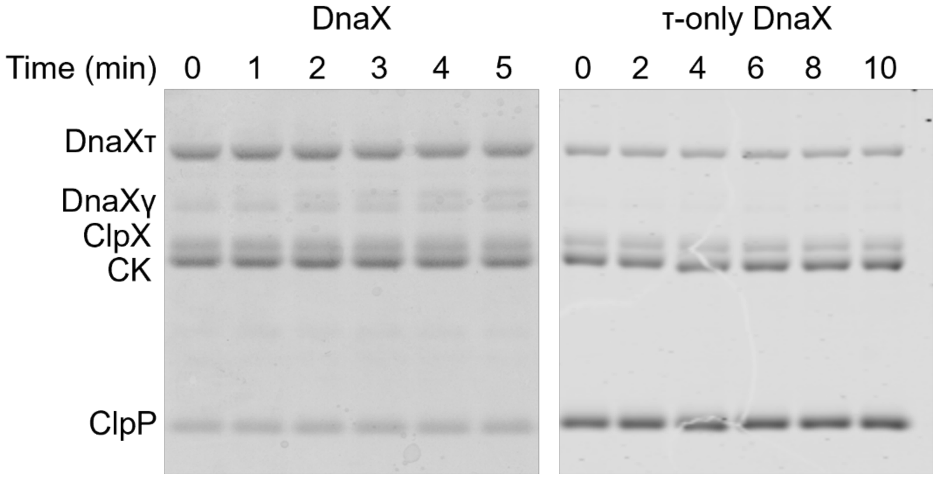


**Supplemental Figure 1. τ-only DnaX is not degraded by ClpXP *in vitro*.** Degradation assay was performed at 30°C as previously described (1) with 2 μM DnaX or τ-only DnaX, 0.2 μM $ClpX_6$, and 0.4 μM $ClpP_{14}$.

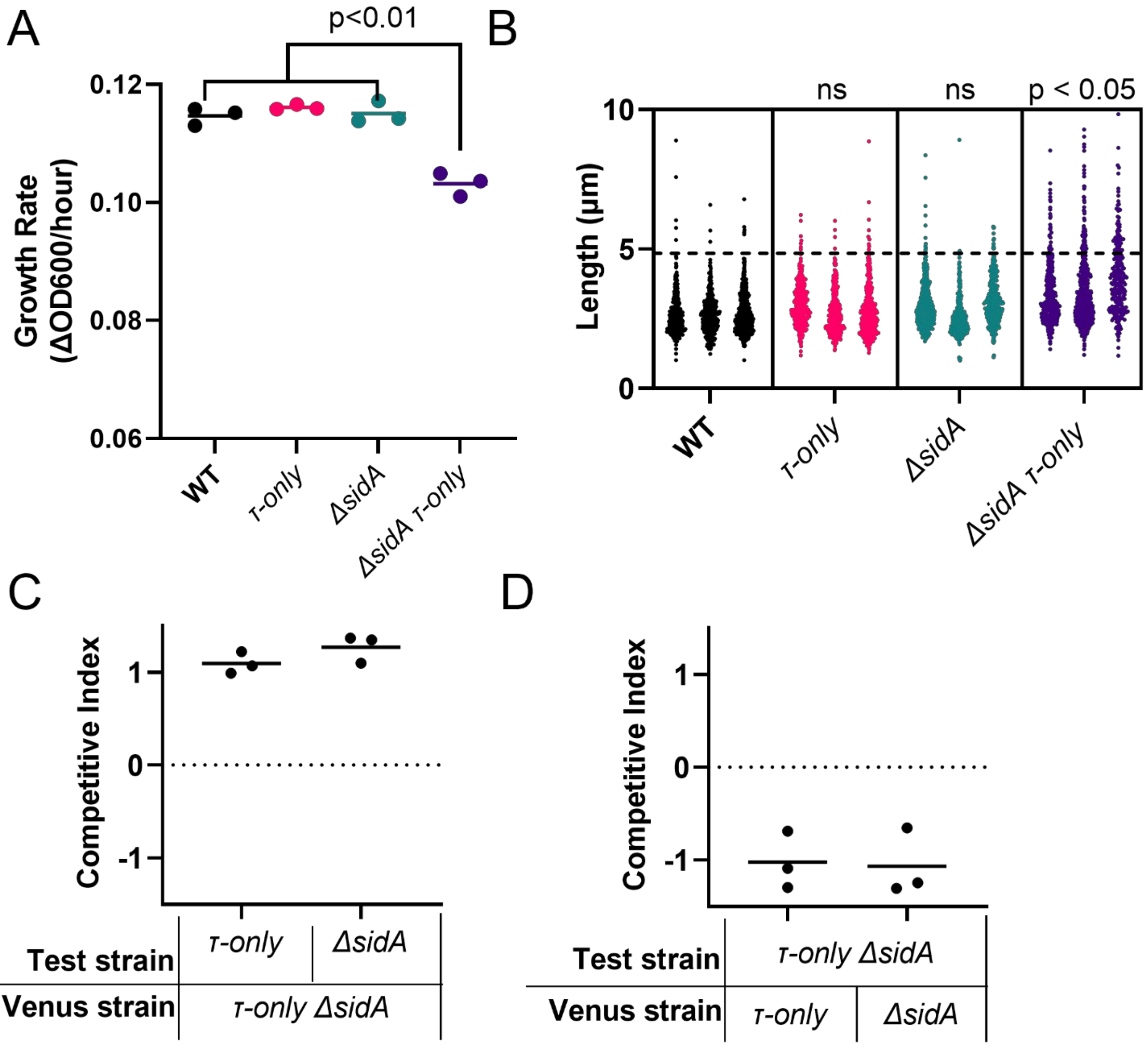


**Supplemental Figure 2. Supplement data corresponding to Figure 4 in main text.** (A) Growth rates of growth curves in Figure 4B. To find growth rate, the slope of the most linear portion of each $\log_{10}$(OD600) vs time graph was take. A two-tailed t-test was used to compare each strain to the wild type. Only the *ΔsidA τ-only* strain was significantly different than the wild type strain. (B) Cell length quantification from Figure 4C. The dashed line represents a benchmark equal to three standard deviations above the mean of the wild type strain cell length (mean = 2.7 μm, benchmark = 4.9 μm). For each strain, the number of cells longer than this benchmark was counted. (Mean ± standard deviation: wild type=4±1, *τ-only*=9±5, *ΔsidA*=9±5, *ΔsidA τ-only*=67±25). A two-tailed t-test was used to compare this value in each strain against the wild-type. Only the *ΔsidA τ-only* strain was significantly different than the wild type strain. ns: not significant. (C) Each single mutant was competed against a Venus-expressing double *ΔsidA τ-only*. Both single mutants outcompete the Venus-expressing *ΔsidA τ-only* strain. (D) The *ΔsidA τ-only* strain was competed against each Venus-expressing single mutant. In both cases, the single mutant outcompeted the *ΔsidA τ-only* strain.

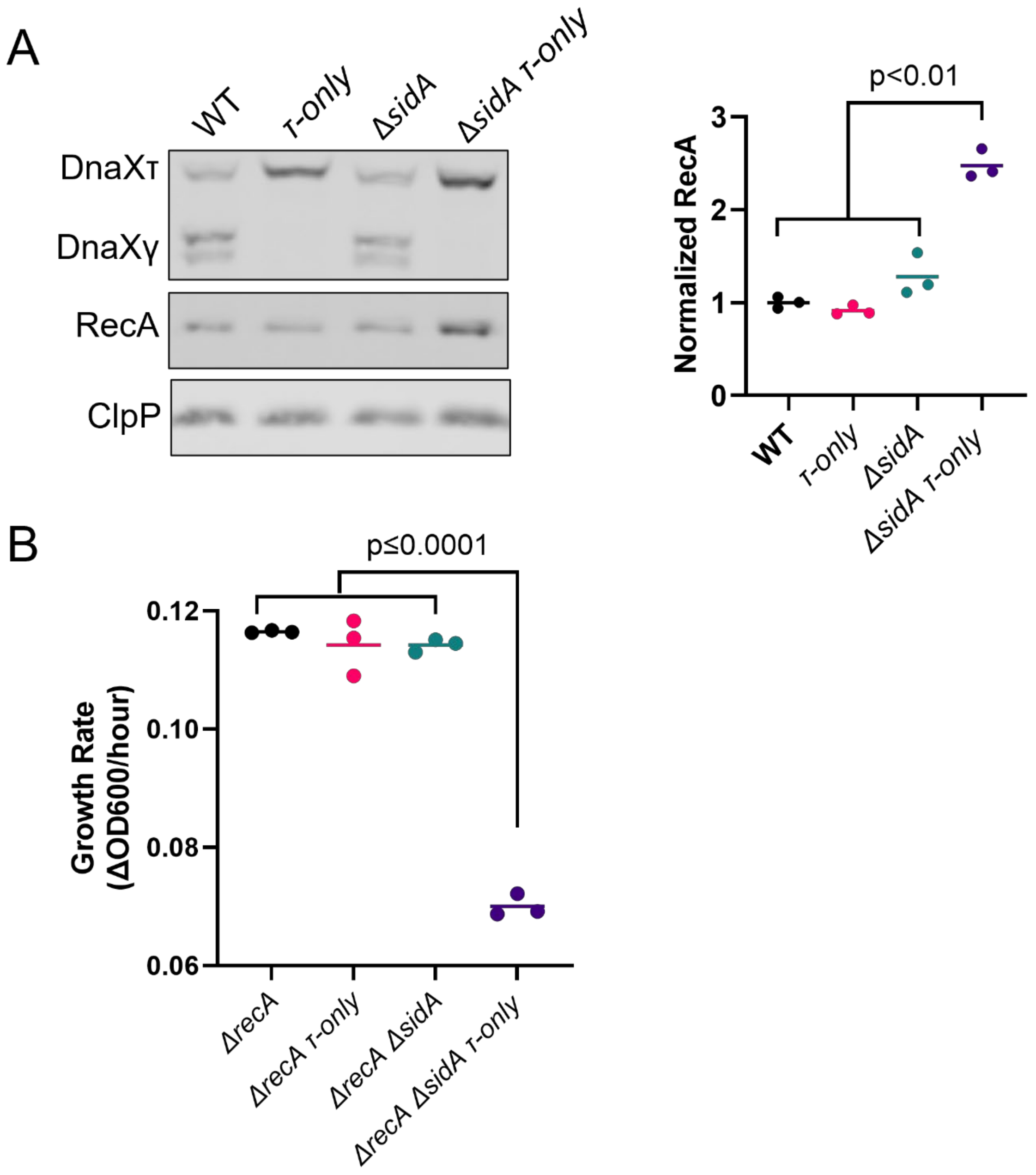


**Supplemental Figure 3. Quantification of RecA western blot and growth rates for growth curves in Figure 5.** (A) Quantification of RecA western blots from Figure 5A. RecA quantification was performed in ImageJ. RecA values were normalized to loading control ClpP, then normalized to the values for the wild type strain. Experiment performed in triplicate, representative images shown. A two-tailed t-test was used to compare each strain to the wild type. Only the *ΔsidA τ-only* strain was significantly different than the wild type strain. (B) Growth rates from growth cures in Figure 5B. Rates determined as in Supplemental Figure 1. A two-tailed t-test was used to compare each strain to the *ΔrecA* strain. Only the *ΔrecA ΔsidA τ-only* strain was significantly different than the *ΔrecA* strain.

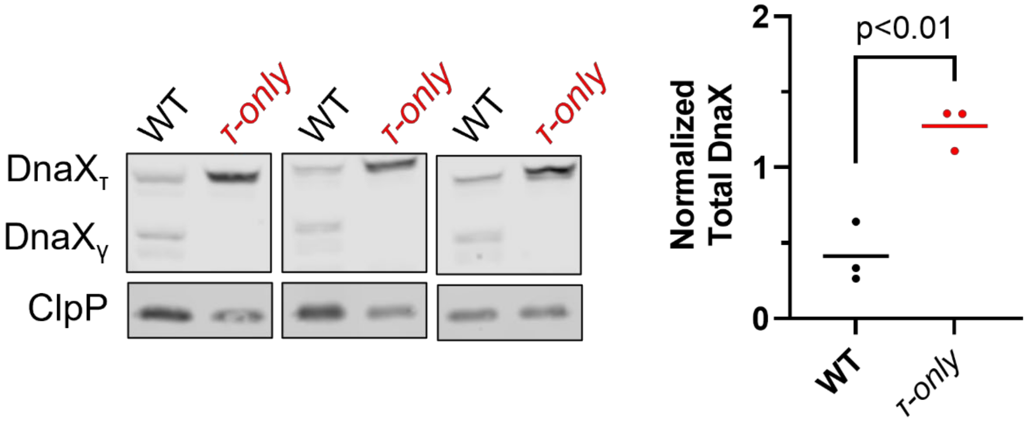


**Supplemental Figure 4. The *τ-only* strain has more total DnaX protein than the wild type strain.** DnaX quantification was performed in ImageJ. Total DnaX values were normalized to loading control ClpP. Two-tailed t-test was performed.

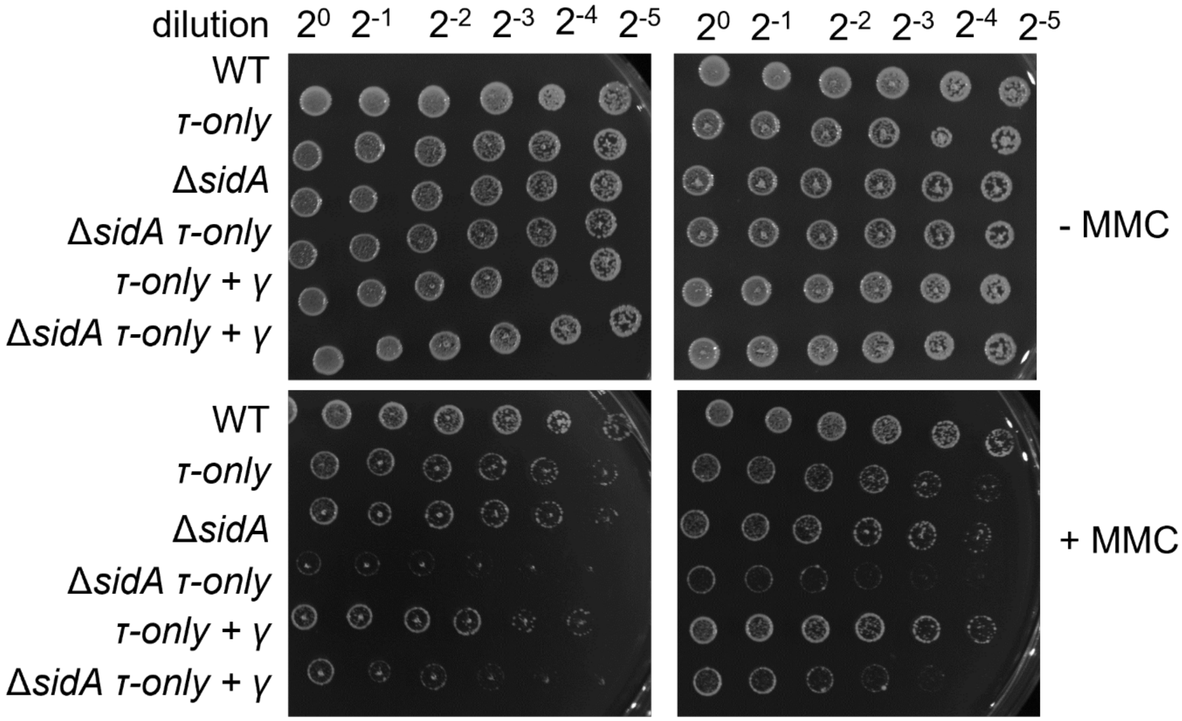


**Supplemental Figure 5. Replicates of mitomycin C sensitivity experiments of Figure 6A.**

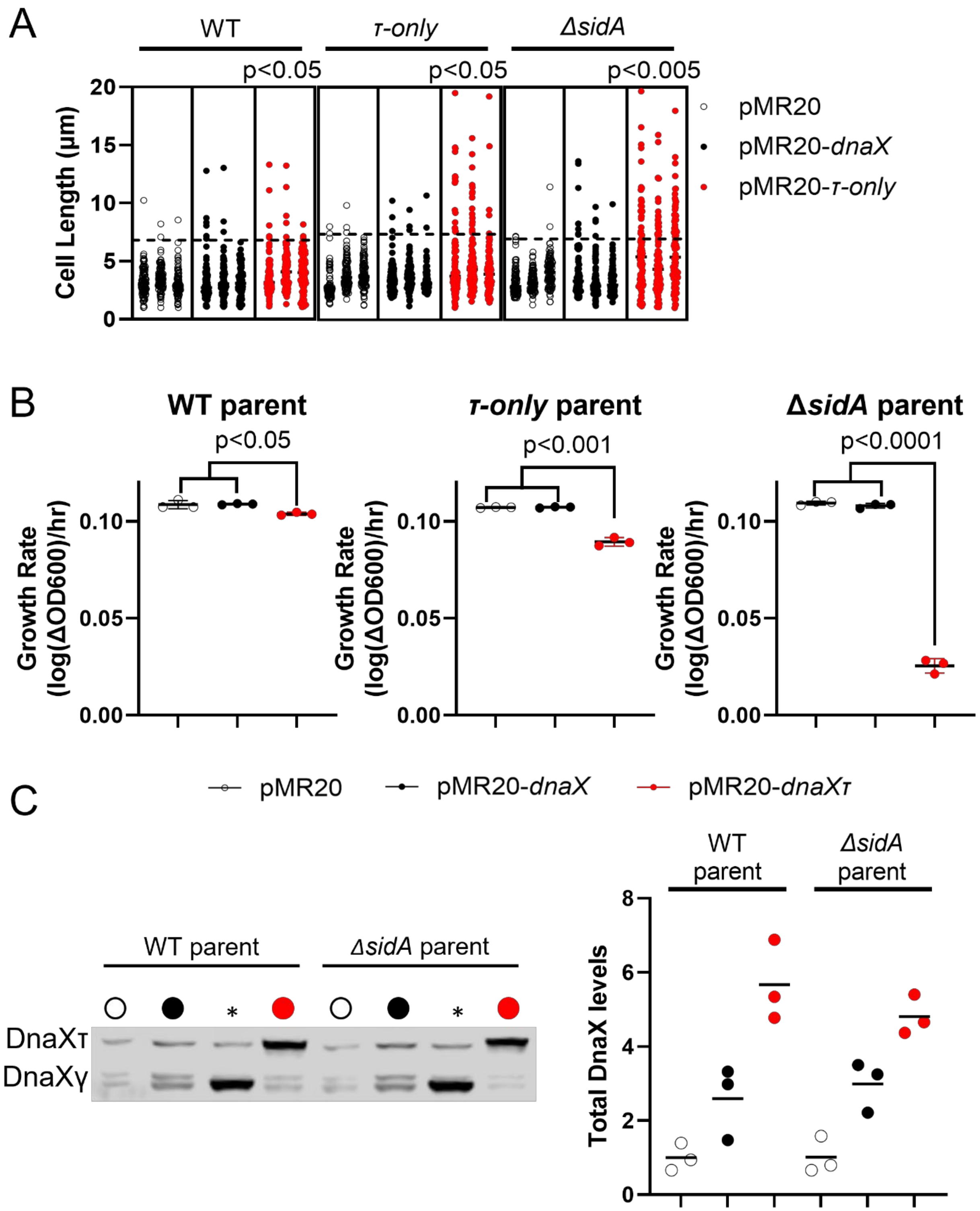
A
WT
τ-only
ΔsidA
p<0.05
p<0.05
p<0.005
Cell Length (µm)
20
15
10
5
0
pMR20
pMR20-dnaX
pMR20-τ-only
B
WT parent
τ-only parent
ΔsidA parent
p<0.05
p<0.001
p<0.0001
Growth Rate (log(ΔOD600)/hr)
0.10
0.05
0.00
C
pMR20
pMR20-dnaX
pMR20-dnaXτ
WT parent
ΔsidA parent
DnaXτ
DnaXγ
*
Total DnaX levels
8
6
4
2
0

38

**Supplemental Figure 6. Supplemental Data corresponding to Figure 7.** Cell length quantification from Figure 7A. For each strain, a benchmark equal to three standard deviations above the mean of the empty vector control was calculated (wild type mean = 3.24 µm, benchmark = 6.81 µm; *τ-only* mean = 2.90 µm, benchmark = 7.36 µm; *ΔsidA* mean = 3.25 µm, benchmark = 6.97 µm). Benchmarks shown as dashed lines. For each plasmid within a parental strain, the number of cells longer than this benchmark was counted. Values shown in Supplemental Table 8. A two-tailed t-test was used to compare this value of the pMR20-*dnaX* and pMR20-*τ-only* strains to the empty vector control. In all three parental strains, the pMR20-*τ-only* strain showed a significant increase from the empty vector, but the pMR20-*dnaX* strain showed little to no effect. (B) Growth rates from growth cures in Figure 7B. Rates determined as in Supplemental Figure 2. Within each parental strain, the pMR20-*dnaX* and pMR20-*τ-only* strain were compared to the empty vector control using the two-tailed t-test. In all cases, the pMR20-*τ-only* strain showed a significant difference from the empty vector control, but the pMR20-*dnaX* did not. (C) Total DnaX in the plasmid-containing wild type and *ΔsidA* strains shows that there is little difference in DnaX expression between parental strain. The pMR20-*dnaX* plasmid increases the total DnaX in the cell by about 2-fold, while the pMR20-*τ-only* plasmid increases total DnaX levels by about 4-fold.
