## Supplemental Table 3 for "Clamp loader processing is important during DNA replication stress"

### τ-only isolate #1
### primary expressed WT DnaX
### final strain expresses τ-only DnaX

***breseq*** version 0.34.1
mutation predictions | marginal predictions | summary statistics | genome diff | command line log

#### Predicted mutations

| evidence | position | mutation | annotation | gene | description |
|---|---|---|---|---|---|
| RA | 283,760 | 2 bp→AC | coding (1631-1632/1827 nt) | *dnaX* → | DNA polymerase III subunit gamma/tau |
| RA | 283,763 | C→A | A545D (GCC→GAC) | *dnaX* → | DNA polymerase III subunit gamma/tau |
| RA | 1,022,865 | +G | coding (999/1203 nt) | *CCNA_00944* → | flagellar hook length determination protein |
| RA | 1,068,007 | Δ1 bp | coding (165/204 nt) | *CCNA_00987* → | stress response protein CsbD |
| JC | 1,682,805 | Δ12 bp | intergenic (+843/-4) | *CCNA_01563* → / → *CCNA_R0142* | 2-dehydro-3-deoxygluconokinase/small non-coding RNA |
| JC | 1,682,979 | +AAGAGAGAGGGA | intergenic (+3/+17) | *CCNA_R0142* → / ← *CCNA_01566* | small non-coding RNA/hypothetical protein |
| RA | 1,891,814 | $(C)_{5\to4}$ | intergenic (-43/-185) | *CCNA_01762* ← / → *CCNA_01763* | YjgP/YjgQ family membrane permease/multifunctional aminopeptidase |
| RA | 1,927,487 | T→C | I189I (ATT→ATC) | *CCNA_01803* → | pyruvate dehydrogenase complex, dihydrolipoamide acetyltransferase component |
| RA | 2,119,025 | G→C | L44V (CTG→GTG) | *CCNA_01973* ← | hypothetical protein |
| RA | 3,026,719 | C→G | intergenic (-4/+71) | *CCNA_02878* ← / ← *CCNA_02879* | hypothetical protein/genetic exchange related protein |
| RA | 3,202,984 | +G | intergenic (-19/+35) | *CCNA_03048* ← / ← *CCNA_03049* | para-aminobenzoate synthetase component I/transporter, MFS superfamily |
| RA | 3,713,704 | C→G | R36P (CGT→CCT) | *CCNA_03557* ← | hypothetical protein |

#### Unassigned missing coverage evidence

| | | | seq id | start | end | size | ←reads | reads→ | gene | description |
|---|---|---|---|---|---|---|---|---|---|---|
| * | * | ÷ | NC_011916 | 1919846 | 1919922 | 77 | 9 [7] | [7] 10 | *CCNA_01792/CCNA_01793* | LSU ribosomal protein L32P/aspartate protease |
| * | * | ÷ | NC_011916 | 2841686 | 2841757 | 72 | 9 [8] | [8] 12 | *CCNA_02687/CCNA_02688* | serine-pyruvate aminotransferase/PuuE-related allantoinase |
| * | * | ÷ | NC_011916 | 2923990 | 2924050 | 61 | 9 [8] | [7] 10 | *CCNA_02761/CCNA_02762* | hypothetical protein/DNA repair protein RadC |

#### Unassigned new junction evidence

| | | seq id | position | reads (cov) | reads (cov) | score | skew | freq | annotation | gene | product |
|---|---|---|---|---|---|---|---|---|---|---|---|
| * | ? | NC_011916 | = 801556 | 23 (0.360) | 27 (0.440) | 23/252 | 0.9 | 55.8% | intergenic (+46/-196) | *CCNA_00743/CCNA_00744* | NnrU family membrane protein/GIY-YIG endonuclease domain protein |
| | ? | NC_011916 | 801695 = | 21 (0.350) | | | | | intergenic (+185/-57) | *CCNA_00743/CCNA_00744* | NnrU family membrane protein/GIY-YIG endonuclease domain protein |
| * | ? | NC_011916 | = 2729001 | 50 (0.780) | 12 (0.200) | 7/248 | 2.9 | 22.1% | intergenic (+35/+197) | *CCNA_02577/CCNA_02578* | phosphoribosylformylglycinamidine synthase, purS component/TetR-family transcriptional regulator |
| | ? | NC_011916 | 2729212 = | 38 (0.630) | | | | | coding (631/645 nt) | *CCNA_02578* | TetR-family transcriptional regulator |
| * | ? | NC_011916 | 3035046 = | 23 (0.360) | 15 (0.310) | 9/202 | 2.0 | 43.2% | intergenic (+60/-170) | *CCNA_02886/CCNA_02887* | RutC-family pyrimidine utilization protein C/hydrolase/acyltransferase family protein |
| | ? | NC_011916 | 3035179 = | 22 (0.450) | | | | | intergenic (+193/-37) | *CCNA_02886/CCNA_02887* | RutC-family pyrimidine utilization protein C/hydrolase/acyltransferase family protein |
| * | ? | NC_011916 | = 3035077 | 25 (0.390) | 6 (0.120) | 5/202 | 2.9 | 22.7% | intergenic (+91/-139) | *CCNA_02886/CCNA_02887* | RutC-family pyrimidine utilization protein C/hydrolase/acyltransferase family protein |
| | ? | NC_011916 | = 3035146 | 22 (0.450) | | | | | intergenic (+160/-70) | *CCNA_02886/CCNA_02887* | RutC-family pyrimidine utilization protein C/hydrolase/acyltransferase family protein |
| * | ? | NC_011916 | = 3037269 | 27 (0.420) | 34 (0.570) | 21/248 | 1.0 | 56.6% | intergenic (+20/+184) | *CCNA_02888/CCNA_02889* | nitrilotriacetate monooxygenase/peptidyl-prolyl cis-trans isomerase |
| | ? | NC_011916 | 3037428 = | 27 (0.450) | | | | | intergenic (+179/+25) | *CCNA_02888/CCNA_02889* | nitrilotriacetate monooxygenase/peptidyl-prolyl cis-trans isomerase |
| * | ? | NC_011916 | = 3061081 | 50 (0.780) | 20 (0.330) | 14/248 | 1.7 | 25.4% | intergenic (+24/-69) | *CCNA_02904/CCNA_02905* | GntR-family transcriptional regulator/very short patch repair (Vsr) endonuclease |
| | ? | NC_011916 | 3061584 = | 71 (1.190) | | | | | intergenic (+57/-37) | *CCNA_02905/CCNA_02906* | very short patch repair (Vsr) endonuclease/acyltransferase family protein |
| * | ? | NC_011916 | 3206047 = | 22 (0.340) | 7 (0.130) | 6/216 | 2.8 | 35.1% | intergenic (-76/+128) | *CCNA_03051/CCNA_03052* | hypothetical protein/acyltransferase |
| | ? | NC_011916 | 3206167 = | 8 (0.150) | | | | | intergenic (-196/+8) | *CCNA_03051/CCNA_03052* | hypothetical protein/acyltransferase |
| * | ? | NC_011916 | = 3317778 | 60 (0.930) | 34 (0.570) | 15/246 | 1.5 | 37.9% | intergenic (+23/+185) | *xylR/argJ* | transcriptional regulator xylR/bifunctional ornithine acetyltransferase/N-acetylglutamate synthase |
| | ? | NC_011916 | 3317943 = | 56 (0.940) | | | | | intergenic (+188/+20) | *xylR/argJ* | transcriptional regulator |

| | | | | | | | | | | | |
|---|---|---|---|---|---|---|---|---|---|---|---|
| | | | | | | | | | | | xylR/bifunctional ornithine acetyltransferase/N-acetylglutamate synthase |
