## Supplemental Table 4 for "Clamp loader processing is important during DNA replication stress"

τ-only isolate #2
primary expressed WT DnaX
final strain expresses τ-only DnaX

***breseq*** version 0.34.1
mutation predictions | marginal predictions | summary statistics | genome diff | command line log

### Predicted mutations

| evidence | position | mutation | annotation | gene | description |
|---|---|---|---|---|---|
| RA | 283,760 | 2 bp→AC | coding (1631-1632/1827 nt) | *dnaX* → | DNA polymerase III subunit gamma/tau |
| RA | 283,763 | C→A | A545D (GCC→GAC) | *dnaX* → | DNA polymerase III subunit gamma/tau |
| RA | 1,022,865 | +G | coding (999/1203 nt) | *CCNA_00944* → | flagellar hook length determination protein |
| RA | 1,068,007 | Δ1 bp | coding (165/204 nt) | *CCNA_00987* → | stress response protein CsbD |
| JC | 1,682,805 | Δ12 bp | intergenic (+843/-4) | *CCNA_01563* → / → *CCNA_R0142* | 2-dehydro-3-deoxygluconokinase/small non-coding RNA |
| JC | 1,682,979 | +AAGAGAGAGGGA | intergenic (+3/+17) | *CCNA_R0142* → / ← *CCNA_01566* | small non-coding RNA/hypothetical protein |
| RA | 1,891,814 | $(C)_{5\rightarrow4}$ | intergenic (-43/-185) | *CCNA_01762* ← / → *CCNA_01763* | YjgP/YjgQ family membrane permease/multifunctional aminopeptidase |
| RA | 1,927,487 | T→C | I189I (ATT→ATC) | *CCNA_01803* → | pyruvate dehydrogenase complex, dihydrolipoamide acetyltransferase component |
| RA | 2,119,025 | G→C | L44V (CTG→GTG) | *CCNA_01973* ← | hypothetical protein |
| RA | 3,026,719 | C→G | intergenic (-4/+71) | *CCNA_02878* ← / ← *CCNA_02879* | hypothetical protein/genetic exchange related protein |
| RA | 3,202,984 | +G | intergenic (-19/+35) | *CCNA_03048* ← / ← *CCNA_03049* | para-aminobenzoate synthetase component I/transporter, MFS superfamily |
| RA | 3,713,704 | C→G | R36P (CGT→CCT) | *CCNA_03557* ← | hypothetical protein |

### Unassigned missing coverage evidence

| | | | seq id | start | end | size | ←reads | reads→ | gene | description |
|---|---|---|---|---|---|---|---|---|---|---|
| * | * | ÷ | NC_011916 | 792397 | 792488 | 92 | 10 [9] | [9] 10 | *[CCNA_00734]–[bamF]* | [CCNA_00734],[bamF] |
| * | * | ÷ | NC_011916 | 2428947 | 2429012 | 66 | 10 [6] | [8] 10 | *CCNA_R0158* | small non-coding RNA |
| * | * | ÷ | NC_011916 | 2923988 | 2924046 | 59 | 10 [8] | [9] 11 | *CCNA_02761/CCNA_02762* | hypothetical protein/DNA repair protein RadC |
| * | * | ÷ | NC_011916 | 3323496 | 3323554 | 59 | 11 [9] | [8] 10 | *CCNA_03164/CCNA_03165* | protein translocase subunit secA/major facilitator superfamily transporter |

### Unassigned new junction evidence

| | | seq id | position | reads (cov) | reads (cov) | score | skew | freq | annotation | gene | product |
|---|---|---|---|---|---|---|---|---|---|---|---|
| * | ? | NC_011916 | = 801556 | 44 (0.570) | 37 (0.510) | 22/254 | 1.1 | 48.1% | intergenic (+46/-196) | *CCNA_00743/CCNA_00744* | NnrU family membrane protein/GIY-YIG endonuclease domain protein |
| | ? | NC_011916 | 801695 = | 38 (0.520) | | | | | intergenic (+185/-57) | *CCNA_00743/CCNA_00744* | NnrU family membrane protein/GIY-YIG endonuclease domain protein |
| * | ? | NC_011916 | = 3037269 | 34 (0.440) | 28 (0.390) | 16/250 | 1.6 | 44.9% | intergenic (+20/+184) | *CCNA_02888/CCNA_02889* | nitrilotriacetate monooxygenase/peptidyl-prolyl cis-trans isomerase |
| | ? | NC_011916 | 3037428 = | 37 (0.510) | | | | | intergenic (+179/+25) | *CCNA_02888/CCNA_02889* | nitrilotriacetate monooxygenase/peptidyl-prolyl cis-trans isomerase |
| * | ? | NC_011916 | = 3061081 | 64 (0.830) | 29 (0.400) | 15/250 | 1.8 | 28.5% | intergenic (+24/-69) | *CCNA_02904/CCNA_02905* | GntR-family transcriptional regulator/very short patch repair (Vsr) endonuclease |
| | ? | NC_011916 | 3061584 = | 86 (1.190) | | | | | intergenic (+57/-37) | *CCNA_02905/CCNA_02906* | very short patch repair (Vsr) endonuclease/acyltransferase family protein |
| * | ? | NC_011916 | = 3317778 | 75 (0.970) | 45 (0.630) | 18/248 | 1.4 | 41.0% | intergenic (+23/+185) | *xylR/argJ* | transcriptional regulator xylR/bifunctional ornithine acetyltransferase/N-acetylglutamate synthase |
| | ? | NC_011916 | 3317943 = | 60 (0.840) | | | | | intergenic (+188/+20) | *xylR/argJ* | transcriptional regulator xylR/bifunctional ornithine acetyltransferase/N-acetylglutamate synthase |
