## Supplemental Table 5 for "Clamp loader processing is important during DNA replication stress"

τ-only isolate #3
primary expressed WT DnaX
final strain expresses τ-only DnaX

***breseq*** version 0.34.1
mutation predictions | marginal predictions | summary statistics | genome diff | command line log

### Predicted mutations

| evidence | position | mutation | annotation | gene | description |
|---|---|---|---|---|---|
| RA | 283,760 | 2 bp→AC | coding (1631-1632/1827 nt) | *dnaX* → | DNA polymerase III subunit gamma/tau |
| RA | 283,763 | C→A | A545D (GCC→GAC) | *dnaX* → | DNA polymerase III subunit gamma/tau |
| RA | 1,022,865 | +G | coding (999/1203 nt) | *CCNA_00944* → | flagellar hook length determination protein |
| RA | 1,068,007 | Δ1 bp | coding (165/204 nt) | *CCNA_00987* → | stress response protein CsbD |
| JC | 1,682,805 | Δ12 bp | intergenic (+843/-4) | *CCNA_01563* → / → *CCNA_R0142* | 2-dehydro-3-deoxygluconokinase/small non-coding RNA |
| JC | 1,682,979 | +AAGAGAGAGGGA | intergenic (+3/+17) | *CCNA_R0142* → / ← *CCNA_01566* | small non-coding RNA/hypothetical protein |
| RA | 1,891,814 | $(C)_{5\rightarrow4}$ | intergenic (-43/-185) | *CCNA_01762* ← / → *CCNA_01763* | YjgP/YjgQ family membrane permease/multifunctional aminopeptidase |
| RA | 1,927,487 | T→C | I189I (ATT→ATC) | *CCNA_01803* → | pyruvate dehydrogenase complex, dihydrolipoamide acetyltransferase component |
| RA | 2,119,025 | G→C | L44V (CTG→GTG) | *CCNA_01973* ← | hypothetical protein |
| RA | 3,026,719 | C→G | intergenic (-4/+71) | *CCNA_02878* ← / ← *CCNA_02879* | hypothetical protein/genetic exchange related protein |
| RA | 3,202,984 | +G | intergenic (-19/+35) | *CCNA_03048* ← / ← *CCNA_03049* | para-aminobenzoate synthetase component I/transporter, MFS superfamily |
| RA | 3,713,704 | C→G | R36P (CGT→CCT) | *CCNA_03557* ← | hypothetical protein |

### Unassigned missing coverage evidence

| | | | seq id | start | end | size | ←reads | reads→ | gene | description |
|---|---|---|---|---|---|---|---|---|---|---|
| * | * | ÷ | NC_011916 | 96476 | 96507 | 32 | 8 [7] | [7] 8 | *CCNA_R0002* | small non-coding RNA |
| * | * | ÷ | NC_011916 | 1315336 | 1315410 | 75 | 8 [6] | [7] 9 | *[CCNA_R0135]* | [CCNA_R0135] |
| * | * | ÷ | NC_011916 | 2924004 | 2924050 | 47 | 8 [7] | [6] 12 | *CCNA_02761/CCNA_02762* | hypothetical protein/DNA repair protein RadC |
| * | * | ÷ | NC_011916 | 3306168 | 3306250 | 83 | 11 [6] | [7] 8 | *CCNA_03150* | hypothetical protein |
| * | * | ÷ | NC_011916 | 3337647 | 3337704 | 58 | 9 [3] | [7] 8 | *CCNA_03177/CCNA_03178* | methylmalonyl-CoA mutase MeaA-like protein/MerR-family transcriptional regulator |

### Unassigned new junction evidence

| | | seq id | position | reads (cov) | reads (cov) | score | skew | freq | annotation | gene | product |
|---|---|---|---|---|---|---|---|---|---|---|---|
| * | ? | NC_011916 | = 801556 | 28 (0.430) | 19 (0.310) | 16/252 | 1.4 | 41.5% | intergenic (+46/-196) | *CCNA_00743/CCNA_00744* | NnrU family membrane protein/GIY-YIG endonuclease domain protein |
| | ? | NC_011916 | 801695 = | 27 (0.440) | | | | | intergenic (+185/-57) | *CCNA_00743/CCNA_00744* | NnrU family membrane protein/GIY-YIG endonuclease domain protein |
| * | ? | NC_011916 | = 3037269 | 30 (0.470) | 27 (0.450) | 14/248 | 1.6 | 48.7% | intergenic (+20/+184) | *CCNA_02888/CCNA_02889* | nitrilotriacetate monooxygenase/peptidyl-prolyl cis-trans isomerase |
| | ? | NC_011916 | 3037428 = | 29 (0.480) | | | | | intergenic (+179/+25) | *CCNA_02888/CCNA_02889* | nitrilotriacetate monooxygenase/peptidyl-prolyl cis-trans isomerase |
| * | ? | NC_011916 | = 3061081 | 43 (0.670) | 33 (0.550) | 22/248 | 0.9 | 37.3% | intergenic (+24/-69) | *CCNA_02904/CCNA_02905* | GntR-family transcriptional regulator/very short patch repair (Vsr) endonuclease |
| | ? | NC_011916 | 3061584 = | 71 (1.180) | | | | | intergenic (+57/-37) | *CCNA_02905/CCNA_02906* | very short patch repair (Vsr) endonuclease/acyltransferase family protein |
| * | ? | NC_011916 | = 3317778 | 59 (0.920) | 27 (0.450) | 18/246 | 1.2 | 33.6% | intergenic (+23/+185) | *xylR/argJ* | transcriptional regulator xylR/bifunctional ornithine acetyltransferase/N-acetylglutamate synthase |
| | ? | NC_011916 | 3317943 = | 52 (0.870) | | | | | intergenic (+188/+20) | *xylR/argJ* | transcriptional regulator xylR/bifunctional ornithine acetyltransferase/N-acetylglutamate synthase |
