## Supplemental Table 6 for "Clamp loader processing is important during DNA replication stress"

**breseq** version 0.34.1
mutation predictions | marginal predictions | summary statistics | genome diff | command line log

τ-only isolate #4
primary expressed WT DnaX
final strain expresses τ-only DnaX

### Predicted mutations

| evidence | position | mutation | annotation | gene | description |
|---|---|---|---|---|---|
| RA | 283,760 | 2 bp→AC | coding (1631-1632/1827 nt) | *dnaX* → | DNA polymerase III subunit gamma/tau |
| RA | 283,763 | C→A | A545D (GCC→GAC) | *dnaX* → | DNA polymerase III subunit gamma/tau |
| RA | 921,646 | Δ1 bp | noncoding (74/108 nt) | *CCNA_R0123* ← | small non-coding RNA |
| RA | 1,022,865 | +G | coding (999/1203 nt) | *CCNA_00944* → | flagellar hook length determination protein |
| RA | 1,068,007 | Δ1 bp | coding (165/204 nt) | *CCNA_00987* → | stress response protein CsbD |
| JC | 1,682,805 | Δ12 bp | intergenic (+843/-4) | *CCNA_01563* → / → *CCNA_R0142* | 2-dehydro-3-deoxygluconokinase/small non-coding RNA |
| JC | 1,682,979 | +AAGAGAGAGGGA | intergenic (+3/+17) | *CCNA_R0142* → / ← *CCNA_01566* | small non-coding RNA/hypothetical protein |
| RA | 1,891,814 | $(C)_{5\rightarrow 4}$ | intergenic (-43/-185) | *CCNA_01762* ← / → *CCNA_01763* | YjgP/YjgQ family membrane permease/multifunctional aminopeptidase |
| RA | 1,927,487 | T→C | I189I (ATT→ATC) | *CCNA_01803* → | pyruvate dehydrogenase complex, dihydrolipoamide acetyltransferase component |
| RA | 2,119,025 | G→C | L44V (CTG→GTG) | *CCNA_01973* ← | hypothetical protein |
| RA | 3,026,719 | C→G | intergenic (-4/+71) | *CCNA_02878* ← / ← *CCNA_02879* | hypothetical protein/genetic exchange related protein |
| RA | 3,202,984 | +G | intergenic (-19/+35) | *CCNA_03048* ← / ← *CCNA_03049* | para-aminobenzoate synthetase component I/transporter, MFS superfamily |
| RA | 3,713,704 | C→G | R36P (CGT→CCT) | *CCNA_03557* ← | hypothetical protein |

### Unassigned missing coverage evidence

| | | | seq id | start | end | size | ←reads | reads→ | gene | description |
|---|---|---|---|---|---|---|---|---|---|---|
| * | * | ÷ | NC_011916 | 2271942 | 2272023 | 82 | 9 [7] | [7] 8 | *CCNA_02121* | ATP-dependent helicase |

### Unassigned new junction evidence

| | | seq id | position | reads (cov) | reads (cov) | score | skew | freq | annotation | gene | product |
|---|---|---|---|---|---|---|---|---|---|---|---|
| * | ? | NC_011916 | = 801556 | 36 (0.540) | 39 (0.620) | 32/254 | 0.4 | 53.8% | intergenic (+46/-196) | *CCNA_00743/CCNA_00744* | NnrU family membrane protein/GIY-YIG endonuclease domai[illegible] protein |
| | ? | NC_011916 | 801695 = | 33 (0.530) | | | | | intergenic (+185/-57) | *CCNA_00743/CCNA_00744* | NnrU family membrane protein/GIY-YIG endonuclease domai[illegible] protein |
| * | ? | NC_011916 | = 1682816 | 0 (0.000) | 10 (0.150) | 9/268 | 2.5 | 100% | intergenic (+854/-4) | *CCNA_01563/CCNA_R0142* | 2-dehydro-3-deoxygluconokinase/sma[illegible] non-coding RNA |
| | ? | NC_011916 | 1682980 = | 0 (0.000) | | | | | intergenic (+4/+16) | *CCNA_R0142/CCNA_01566* | small non-coding RNA/hypothetical protein |
| * | ? | NC_011916 | = 2728989 | 80 (1.210) | 16 (0.260) | 9/248 | 2.4 | 23.2% | intergenic (+23/+209) | *CCNA_02577/CCNA_02578* | phosphoribosylformylglycinamidine synthase, purS component/TetR-family transcriptional regulator |
| | ? | NC_011916 | 2729200 = | 32 (0.520) | | | | | coding (643/645 nt) | *CCNA_02578* | TetR-family transcriptional regulator |
| * | ? | NC_011916 | = 3037269 | 29 (0.440) | 31 (0.500) | 17/250 | 1.3 | 51.6% | intergenic (+20/+184) | *CCNA_02888/CCNA_02889* | nitrilotriacetate monooxygenase/peptidyl-prolyl cis-trans isomerase |
| | ? | NC_011916 | 3037428 = | 31 (0.500) | | | | | intergenic (+179/+25) | *CCNA_02888/CCNA_02889* | nitrilotriacetate monooxygenase/peptidyl-prolyl cis-trans isomerase |
| * | ? | NC_011916 | = 3061081 | 41 (0.620) | 24 (0.390) | 13/250 | 1.7 | 25.6% | intergenic (+24/-69) | *CCNA_02904/CCNA_02905* | GntR-family transcriptional regulator/very short patch repair (Vsr) endonuclease |
| | ? | NC_011916 | 3061584 = | 101 (1.640) | | | | | intergenic (+57/-37) | *CCNA_02905/CCNA_02906* | very short patch repair (Vsr) endonuclease/acyltransferase family protein |
| * | ? | NC_011916 | = 3317791 | 44 (0.660) | 42 (0.690) | 18/248 | 1.2 | 38.6% | intergenic (+36/+172) | *xylR/argJ* | transcriptional regulator xylR/bifunctional ornithine acetyltransferase/N-acetylglutamate synthase |
| | ? | NC_011916 | 3317956 = | 93 (1.520) | | | | | intergenic (+201/+7) | *xylR/argJ* | transcriptional regulator xylR/bifunctional ornithine acetyltransferase/N-acetylglutamate synthase |
