## supplemental Table 7 for "Clamp loader processing is important during DNA replication stress"

τ-only isolate #5
primary expressed τ-only DnaX
final strain expresses τ-only DnaX

***breseq*** version 0.34.1
mutation predictions | marginal predictions | summary statistics | genome diff | command line log

### Predicted mutations

| evidence | position | mutation | annotation | gene | description |
|---|---|---|---|---|---|
| RA | 283,760 | 2 bp→AC | coding (1631-1632/1827 nt) | *dnaX* → | DNA polymerase III subunit gamma/tau |
| RA | 283,763 | C→A | A545D (GCC→GAC) | *dnaX* → | DNA polymerase III subunit gamma/tau |
| RA | 921,646 | Δ1 bp | noncoding (74/108 nt) | *CCNA_R0123* ← | small non-coding RNA |
| RA | 1,022,865 | +G | coding (999/1203 nt) | *CCNA_00944* → | flagellar hook length determination protein |
| RA | 1,068,007 | Δ1 bp | coding (165/204 nt) | *CCNA_00987* → | stress response protein CsbD |
| JC | 1,682,805 | Δ12 bp | intergenic (+843/-4) | *CCNA_01563* → / → *CCNA_R0142* | 2-dehydro-3-deoxygluconokinase/small non-coding RNA |
| JC | 1,682,979 | +AAGAGAGAGGGA | intergenic (+3/+17) | *CCNA_R0142* → / ← *CCNA_01566* | small non-coding RNA/hypothetical protein |
| RA | 1,891,814 | (C)$_{5\rightarrow4}$ | intergenic (-43/-185) | *CCNA_01762* ← / → *CCNA_01763* | YjgP/YjgQ family membrane permease/multifunctional aminopeptidase |
| RA | 1,927,487 | T→C | I189I (ATT→ATC) | *CCNA_01803* → | pyruvate dehydrogenase complex, dihydrolipoamide acetyltransferase component |
| RA | 2,119,025 | G→C | L44V (CTG→GTG) | *CCNA_01973* ← | hypothetical protein |
| RA | 3,026,719 | C→G | intergenic (-4/+71) | *CCNA_02878* ← / ← *CCNA_02879* | hypothetical protein/genetic exchange related protein |
| RA | 3,030,603 | C→G | intergenic (+147/+63) | *CCNA_02881* → / ← *CCNA_02882* | oxoglutarate semialdehyde dehydrogenase/cbb3-type cytochrome C oxidase, subunit III |
| RA | 3,202,984 | +G | intergenic (-19/+35) | *CCNA_03048* ← / ← *CCNA_03049* | para-aminobenzoate synthetase component I/transporter, MFS superfamily |
| RA | 3,713,704 | C→G | R36P (CGT→CCT) | *CCNA_03557* ← | hypothetical protein |

### Unassigned missing coverage evidence

| | | | seq id | start | end | size | ←reads | reads→ | gene | description |
|---|---|---|---|---|---|---|---|---|---|---|
| * | * | ÷ | NC_011916 | 1137201 | 1137245 | 45 | 9 [4] | [2] 17 | *CCNA_R0130/CCNA_01042* | small non-coding RNA/TonB-dependent receptor |
| * | * | ÷ | NC_011916 | 1919843 | 1919927 | 85 | 9 [8] | [6] 12 | *CCNA_01792/CCNA_01793* | LSU ribosomal protein L32P/aspartate protease |
| * | * | ÷ | NC_011916 | 2428977 | 2429039 | 63 | 9 [8] | [7] 10 | *[CCNA_R0158]* | [CCNA_R0158] |
| * | * | ÷ | NC_011916 | 2923980 | 2924047 | 68 | 9 [8] | [8] 9 | *CCNA_02761/CCNA_02762* | hypothetical protein/DNA repair protein RadC |

### Unassigned new junction evidence

| | | seq id | position | reads (cov) | reads (cov) | score | skew | freq | annotation | gene | product |
|---|---|---|---|---|---|---|---|---|---|---|---|
| * | ? | NC_011916 | = 801556 | 27 (0.420) | 29 (0.480) | 18/250 | 1.3 | 49.3% | intergenic (+46/-196) | *CCNA_00743/CCNA_00744* | NnrU family membrane protein/GIY-YIG endonuclease domain protein |
| | ? | NC_011916 | 801695 = | 34 (0.560) | | | | | intergenic (+185/-57) | *CCNA_00743/CCNA_00744* | NnrU family membrane protein/GIY-YIG endonuclease domain protein |
| * | ? | NC_011916 | = 3037269 | 38 (0.600) | 23 (0.390) | 14/246 | 1.7 | 43.6% | intergenic (+20/+184) | *CCNA_02888/CCNA_02889* | nitrilotriacetate monooxygenase/peptidyl-prolyl cis-trans isomerase |
| | ? | NC_011916 | 3037428 = | 24 (0.400) | | | | | intergenic (+179/+25) | *CCNA_02888/CCNA_02889* | nitrilotriacetate monooxygenase/peptidyl-prolyl cis-trans isomerase |
| * | ? | NC_011916 | = 3061081 | 59 (0.930) | 18 (0.300) | 11/246 | 2.1 | 21.6% | intergenic (+24/-69) | *CCNA_02904/CCNA_02905* | GntR-family transcriptional regulator/very short patch repair (Vsr) endonuclease |
| | ? | NC_011916 | 3061584 = | 76 (1.280) | | | | | intergenic (+57/-37) | *CCNA_02905/CCNA_02906* | very short patch repair (Vsr) endonuclease/acyltransferase family protein |
| * | ? | NC_011916 | = 3317791 | 31 (0.490) | 23 (0.390) | 14/244 | 1.7 | 29.9% | intergenic (+36/+172) | *xylR/argJ* | transcriptional regulator xylR/bifunctional ornithine acetyltransferase/N-acetylglutamate synthase |
| | ? | NC_011916 | 3317956 = | 79 (1.340) | | | | | intergenic (+201/+7) | *xylR/argJ* | transcriptional regulator xylR/bifunctional ornithine acetyltransferase/N-acetylglutamate synthase |
